# Sensorimotor encoding of epistemic value during goal-directed causal learning

**DOI:** 10.64898/2026.06.03.729806

**Authors:** Ruggero Basanisi, Emmanuel Daucé, Etienne Combrisson, Mehdi Khamassi, Mateus Joffily, Andrea Brovelli

## Abstract

Understanding the neural and computational mechanisms underlying goal-directed causal learning is a central challenge in both cognitive neuroscience and artificial intelligence. This cognitive function depends on balancing reward maximization with information seeking. Although substantial progress has been made in characterizing the neural basis of reward-driven learning, it remains unclear whether and how intrinsic informational value is represented in the brain and propagated through cortico-cortical interactions. Here, we dissociate information-seeking from reward maximisation during goal-directed causal learning using a novel behavioural paradigm in which participants estimate action–outcome contingencies without extrinsic incentives. Bayesian computational modelling reveals that while random exploration dominates behaviour, expected information gain (EIG) explains a significant proportion of exploratory choices in a subset of sessions. Using magnetoencephalography combined with information-theoretic analyses of high-gamma activity, we show that EIG is encoded in the left sensorimotor cortex at the time of action execution, specifically in dorsal premotor and primary motor cortex, and broadcast to primary somatosensory cortex. These findings are consistent with Active Inference predictions that EIG constitutes a key computational drive for exploratory action selection. However, they challenge the view that epistemic valuation exclusively recruits prefrontal reward circuits, and instead they support an embodied account in which the premotor and sensorimotor system mediates the intrinsic valuation during first-personal interventional causal learning.

## INTRODUCTION

A fundamental facet of human cognition is the ability to learn about the causal structure of the world. Human causal reasoning can be characterized as the process by which statistical regularities derived from sensory observations and behavioral interventions are transformed into structured representations of cause–effect relationships (Goddu and Gopnik 2024). It allows us to climb “the ladder of causation” (Pearl and Mackenzie 2018), moving from purely statistical associations between variables (first level or ladder of causation: association) to causal beliefs derived from behavioral manipulations (second level: intervention, “what happens if we double the price?”), and finally to counterfactual reasoning (third level, “what would have happened if I had acted differently?”), which requires retrospective inference. Causal learning is fundamental to understanding human cognition and remains a central topic in contemporary Artificial Intelligence research. Despite notable advances, standard large language model (LLM) agents remain limited in their capacity to acquire causal understanding and to construct coherent models of the world (Pezzulo et al. 2024; Khamassi et al. 2024). Indeed, causal reasoning currently represents an open challenge in frontier Artificial Intelligence research, with emerging architectures implementing structured, interventional, and counterfactual reasoning to varying degrees (Hafner et al. 2025). Understanding the neural and computational foundations of human causal learning is therefore critical for both neuroscience and potential applications in AI.

We investigated a central form of causal learning, which supports the ability to learn beliefs about the consequences of our actions. This ability provides the basis for adaptive human agency, rational decision-making and, in general, allows people to adapt to changing environments, engage in meaningful life and social interactions all along their life (Bandura 1997). Learning the causal relation between actions and outcomes is supported by the so-called goal-directed system (Tolman 1948; Dickinson 1994; Dolan and Dayan 2013; Morris et al. 2022). Goal-directed behaviours are thought to emerge from the coordinated activity of neural populations distributed over the associative fronto-striatal circuit (Haber and Knutson 2009) and the limbic “reward” system (Haber and Knutson 2009). More specifically, goal-directed learning recruits cortical circuits including the lateral and medial prefrontal cortex (lPFC and mPFC), posterior parietal cortex, orbito-frontal cortex (OFC), temporal gyrus and visual cortex, and the temporal parietal junction (Liljeholm et al. 2011; Tanaka et al. 2008; Liljeholm et al. 2013; Norton and Liljeholm 2020; Jocham et al. 2016; Walton et al. 2010; Morris et al. 2022; Averbeck and Costa 2017; Averbeck and O’Doherty 2021; Bartolo and Averbeck 2020). Goal-directed causal learning is rooted in the balance between reward and information maximisation (Cohen et al. 2007; Schwartenbeck et al. 2019; Gottlieb et al. 2013; Cockburn et al. 2022; Badre et al. 2012; Frank et al. 2009; Mehlhorn et al. 2015).

On the one hand, a key determinant of action valuation are (extrinsic) reward prediction errors (RPEs), which signal whether an outcome is better or worse than expected. Action values carrying information about the expected future utility are carried and updated in the orbitofrontal cortex (OFC) and ventromedial prefrontal cortex (vmPFC) (Daw and O’Doherty 2014; Rangel et al. 2008; Padoa-Schioppa and Assad 2006; Rushworth et al. 2011). Lateral prefrontal regions then contribute to comparing candidate actions and implementing action plans to mediate value-based decision making and action selection (Morris et al. 2014; Cai and Padoa-Schioppa 2014; Miller and Cohen 2001). On the other hand, information maximization provides an additional learning objective supporting novelty seeking and exploration (Kidd and Hayden 2015; Modirshanechi et al. 2025; Gottlieb et al. 2013), and they have been proposed to play a key role in curiosity-driven learning (Schwartenbeck et al. 2019; Gottlieb and Oudeyer 2018). At the neural level, information prediction errors, reflecting the difference between obtained and expected information gain, is encoded by the activity of dopaminergic neurons (Bromberg-Martin and Hikosaka 2009; Bromberg-Martin et al. 2024) and recruit the mesolimbic reward circuitry in humans (Charpentier et al. 2018). Neural correlates of subjective value of information in instrumental settings (i.e., information that can be used to guide future actions and future outcomes) have also been observed in a distributed network including the ventral striatum, the vmPFC, the middle and superior frontal gyrus (i.e., the dorsolateral prefrontal cortex dlPFC), and posterior cingulate cortex (Kobayashi and Hsu 2019; Kobayashi et al. 2021; Kobayashi and Kable 2024). At the large-scale level, information gain has been mapped to the activity of distributed brain networks including the middle frontal gyrus, the insula and the intraparietal sulcus (Fouragnan et al. 2018; Liakoni et al. 2022; Modirshanechi et al. 2023; Gläscher et al. 2010; Lee et al. 2014). More recently, information gain has been shown to be contained via synergistic and higher-order (i.e., beyond pairwise relations) functional interactions and broadcast to prefrontal reward circuits during goal-directed learning (Combrisson et al. 2025). Nevertheless, the neural circuitry supporting how reward and information values are integrated to drive action selection during causal learning remains poorly understood.

From the theoretical point of view, information seeking and exploration are formalised using model-based reinforcement algorithms or Bayesian models (Friston et al. 2015, 2017; Faraji et al. 2018; Gläscher et al. 2010; Liakoni et al. 2021; Yu and Dayan 2005; Modirshanechi et al. 2022). Information gain (IG) is normally formalised in terms of Bayesian surprise (Baldi and Itti 2010; Itti and Baldi 2009), which quantifies how much the agent’s belief changes given new observation. The value of actions is not solely determined by the expected utility (or extrinsic value), but also by the informative (intrinsic, epistemic, or non-instrumental) value that is expected to be gained, which helps reduce uncertainty about environmental contingencies (Cohen et al. 2007; White et al. 2019; Bromberg-Martin and Hikosaka 2009; Friston et al. 2015; Tishby and Polani 2011). A normative model for a unified action-perception cycle and accounting for planning and learning and combining reward maximization with information gain mechanisms is provided by Active Inference (Parr et al. 2022). Within this framework, actions are thought to be selected to maximise expected free energy, which is equal to the sum of extrinsic (expected utility) and epistemic (expected information gain, EIG) values (Friston et al. 2015).

Despite recent progress in the theoretical and behavioral underpinnings, it remains unclear whether and how intrinsic values based on EIG are encoded in the brain and broadcast via cortico-cortical interactions. To address this question, we designed a novel experimental task that isolates information seeking from reward maximisation processes during the acquisition of causal relations between actions and outcomes. Behavioral patterns were analysed using a Bayesian computational modelling approach. The inference of action-outcomes contingencies during learning was modelled by means of an ideal Bayesian observer. The action selection process was modelled using EIG as an intrinsic valuation signal. Although random exploration was found to be the dominant behavioral determinant, we found that EIG explains behavior better than a random agent in a significant portion of learning sessions, suggesting it helps shape strategies during action–outcome causal learning. We then investigated whether EIG is contained and broadcast in brain circuits by means of information theoretical analyses of source-level high-gamma activity (HGA, from 60 to 120Hz) from magnetoencephalography (MEG). We observed that EIG correlates with HGA from the left sensorimotor regions, including the motor and dorsal premotor regions, and the somatosensory cortex, at the time of motor response. We then performed Feature-specific Information Transfer (FIT) analyses between HGA time series (Celotto et al. 2023) and we observed that EIG was first present in the motor cortex and then broadcast to the somatosensory regions. Overall, our results suggest that EIG is a key intrinsic value signal that shapes strategies during goal-directed causal learning and is implemented in the motor system. These results align with embodied cognition accounts, in which cognitive processes are grounded in sensorimotor systems.

## RESULTS

### Goal-directed learning task and the acquisition of causal action-outcome contingencies

We designed a novel task to dissociate reward maximization from information-seeking by isolating learning about action–outcome causality in the absence of explicit reward incentives. Participants performed a self-paced decision task (referred to here as the “volleyball task”), in which they evaluated the causal impact of a new player on a volleyball team using a simulated environment. On each trial, participants chose whether to include (“Play”) or not (“Not Play”) the new player in their team for a fictitious match. After a fixed delay of 0.3s, the outcome of the match (win or loss) was presented (Fig. 1A). Across 40 trials per block (representing 40 matches of a sport season), participants sampled action–outcome contingencies and, at the end of each block, were required to report a continuous causal judgment (ranging from −100 to 100) reflecting the inferred causal effect of the player on team performance. Positive causal scores reflect performance improvement, with +100 corresponding to a player whose inclusion always results in a win, whereas negative scores indicate performance deterioration, with −100 corresponding to a player whose inclusion always results in a loss. A score of 0 would correspond to a player with null causal effect on the result of the match, carrying no evident changes in team performance. The causal relation was manipulated via the contingency value ΔP, computed as P(Win|Play) − P(Win|Not Play), with five levels spanning negative, null, and positive effects (−0.6, −0.3, 0, 0.3, 0.6). Each ΔP level was instantiated by three probability combinations, yielding 15 independent learning blocks, representing 15 different players in 15 different seasons, presented in randomized order (Fig. 1B). This design enabled the examination of exploratory behavior and causal inference under controlled statistical contingencies, without confounding reward maximization.

**Figure 1.**
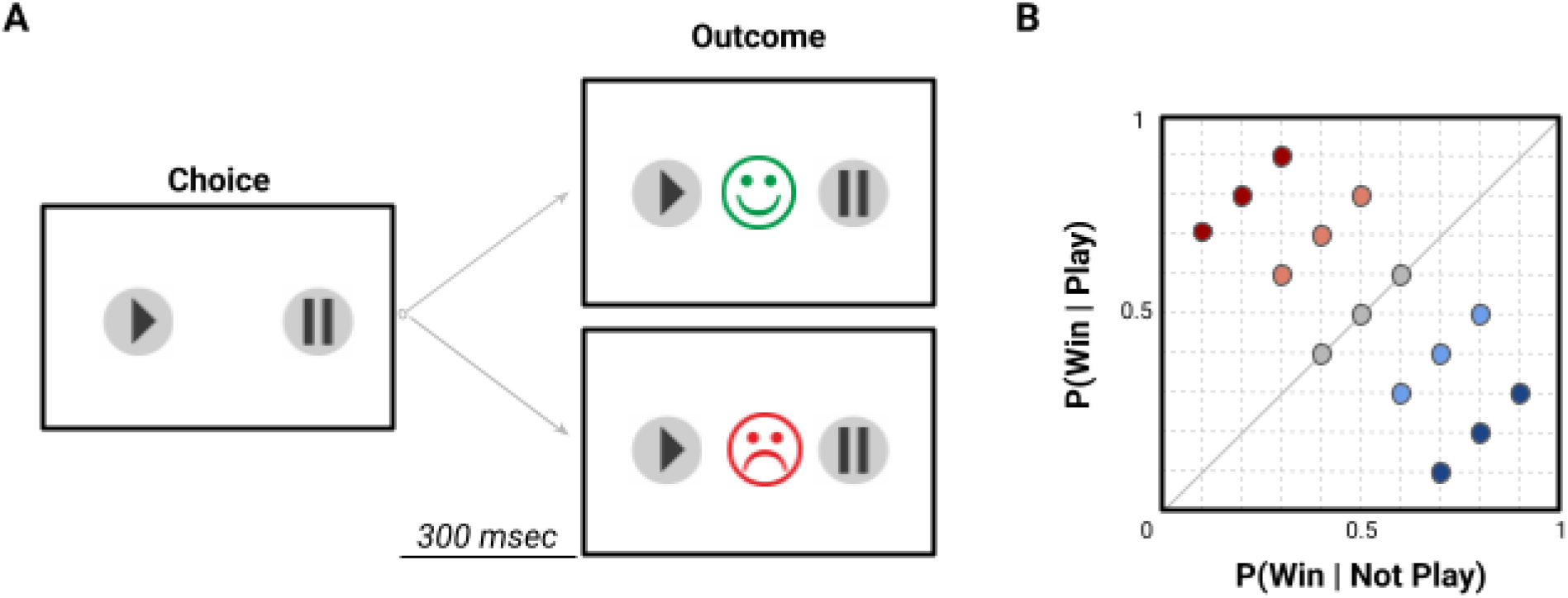
Experimental design and contingency table. **(A)** On each trial, participants could choose to simulate a match either with or without the new player by clicking on one out of two buttons, associated with Play (arrow) and NotPlay (vertical bars; “Pause” symbol) icons, respectively. The position of the Play and NotPlay button were kept constant in each learning block, and they were randomised across blocks. After a fixed delay of 0.3s the outcome image informed the participants whether the fictive match was won or lost by the team. The outcome image was presented for 1.5s. The disappearance of the outcome image instructed the participants that another match could be simulated at any time. **(B)** Contingency plot showing 15 combinations of conditional probabilities. Colors indicate the contingency ΔP equal to 0.6 (dark red contingency), 0.3 (pink), 0 (grey), 0.3 (light blue) and 0.6 (blue).

In order to investigate whether participants acquired the action-outcome causal scores, we made the assumption that causal judgments are grounded in the statistical relationship between potential causes and their outcomes. Central to these approaches is the Δ*P* model (Jenkins and Ward 1965; “Website,” n.d.; Allan et al. 2008; Morris et al. 2022), which defines causal strength as the difference between the probability of the effect *E* when the cause *C* is present and when it is absent (indicated by ¬):

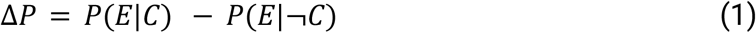

When the causal relation concerns actions and outcomes, Δ*P* is referred to as the action–outcome contingency. Its value ranges from –1 to +1 and correlates with subjective judgments of instrumental causality. A positive Δ*P* leads to the impression that one’s action causes the outcome, whereas a negative Δ*P* supports the belief that the action prevents the outcome. When Δ*P* is close to zero, individuals typically perceive no causal relation between action and outcome (Shanks and Dickinson 1991).

To model the trial-by-trial formation of causal beliefs, we implemented an ideal Bayesian observer based on conjugate Beta-Binomial inference (see Methods). Subjective causal scores were operationalised as the contingency ΔP = P(Win|Play) − P(Win|¬Play), with each conditional win probability estimated independently. Match outcomes were treated as Bernoulli trials, and the posterior distribution of the conditional probabilities after observing *k* successes in *n* trials updates analytically to Beta(α_0_ + k, β_0_ + n − k), with posterior mean (α_0_ + k)/(α_0_ + β_0_ + n). The Δ*P* estimate at each trial was computed as the difference between the posterior means for the Play and Not-Play choices. This formulation yields a computationally tractable, normatively grounded account of incremental causal learning from binary feedback. Figure 2A shows the mean evolution of Δ*P* estimated for the five levels of causal effect, assuming an ideal Bayesian observer. In order to test the accuracy of the reported causal scores and ideal observed assumption, we computed the linear correlations between the subjective scores and the Δ*P* estimated at the end of learning (trial 40). Figure 2B shows a significant linear correlation (R^2 = 0.85, Pearson r = 0.92, p-value < 0.001), indicating that the acquisition of causal action-outcome relations in humans parallels a learning update predicted by the ideal Bayesian observer model. Finally, we investigated the behavioral patterns during learning. Given that the aim of the task was to provide an accurate causal score rather than to have the team win matches, we predicted that an information-seeking, rather than a reward-maximisation, pattern would have emerged. In the literature, information seeking and exploration in humans has been shown to be based on a mixture of random and directed strategies (Wilson et al. 2014; Gershman 2018). Random exploration is a deviation from the most rewarding option, whereas directed exploration selectively samples options that are informative (information seeking), that are associated with the highest uncertainty. We expected participants to make choices to maximise information gain, and randomly sample Play and Not-Play. Indeed, the probability of Play was approximately 50% for all trials during learning and across contingency values (Fig. 2C). Overall, these results indicate that participants predominantly adopted an information seeking and explorative behavior for the acquisition of causal action-outcome relations, using a near optimal update of internal beliefs as predicted by the ideal Bayesian observer model.

**Figure 2.**
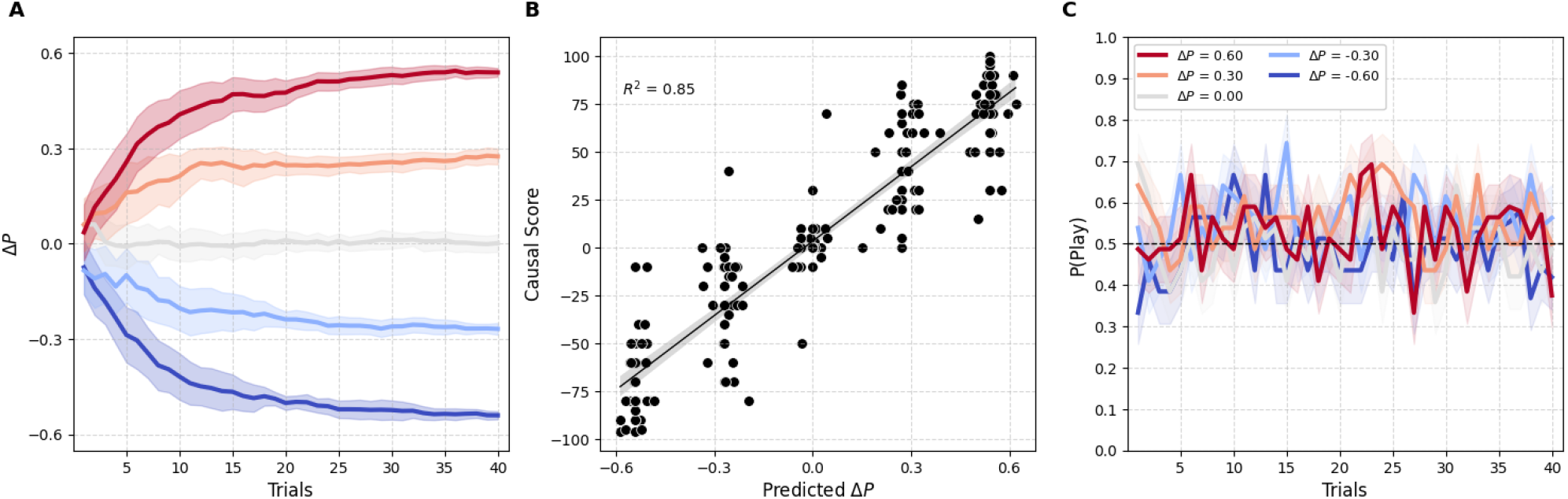
Causal scores and behavioral strategies. **(A)** Evolution of the ΔP estimated for the five levels of causal effect, assuming an ideal Bayesian observer. **(B)** Correlation between the subjects’ causal scores and the ΔP predicted by the model at the end of the learning session (trial 40), assuming an ideal Bayesian observer. **(C)** Mean probability to choose “Play” as a function of learning trials and ΔP. Participants choose Play with a probability of approximately 0.5.

### Complementary action valuation systems and information seeking behavioral strategies

We then investigated behavioural choices with the aim of dissociating contributions from random and directed exploration during learning. Random exploration can be simply defined as a behavioral strategy choosing Play with probability equal to 0.5. However, directed exploration is classically defined as selectively sampling options that are informative, that are associated with the maximum expected information gain (EIG). This notion aligns with predictions from Active Inference (AI), which represents a normative model of perception, action and planning in terms of variational free energy minimization (Parr et al. 2022). In the context of goal-directed learning and planning, AI predicts that agents select actions to minimize the expected free energy of future outcomes. Since the negative free energy (or quality of a policy) can be decomposed into extrinsic and epistemic (or intrinsic) value, minimizing expected free energy is equivalent to maximizing extrinsic value (or expected utility, defined in terms of prior preferences or goals), while maximizing information gain or intrinsic value (or reducing uncertainty about the causes of valuable outcomes or epistemic value) (Friston et al. 2015).

In order to quantify the contribution of random versus directed exploration based on EIG, we took, as a baseline model, an agent that samples each action randomly with no learning and no valuation systems. We then modelled the probability of performing an action as a weighted mixture of two components: a random baseline and a general value-driven component (see Methods). The mixing weight *w* quantifies, for each individual and session, how much of their behaviour is governed by structured value signals relative to random exploration: *w* = 0 corresponds to pure random behaviour while *w* = 1 corresponds to fully deterministic, value-driven choices. We first tested EIG as a main value-driven component, which reflected the expected reduction in uncertainty about the environment regardless of reward value. We observed that the distribution of weights was strongly skewed towards zero, indicating that most of the behavioral strategies were dominated by random exploration. However, a large portion of sessions were characterised by a high contribution of EIG, associated with values equal or larger than 0.5. As a reminder, a weight of 0.5 indicates a comparable contribution of random and EIG-based policy (Fig. 3A, blue dots and distribution). As a control analysis, we investigated whether the subjective expected utility (EU) of actions, defined as the probability-weighted sum of log-softmax utilities over binary outcomes (see Methods), could influence behavioral choices. We fitted the mixing weight by assuming a random agent influenced by EU, and we observed that the number of sessions displaying a comparable contribution of random and EU-based choices (i.e., weight equal or larger than 0.5) was rare (Fig. 3A, orange circles and distribution). Finally, we tested whether expected free energy (EFE), equal to the sum of EIG and EU as proposed by Active Inference, would drive behaviour. Since EFE is highly correlated with EIG in the current task, we expected the weight to resemble the distribution of values for EIG. Indeed, a significant portion of sessions displayed weights larger than 0.5 (Fig. 3A, green circles and distribution) indicating a strong contribution of EFE.

**Figure 3.**
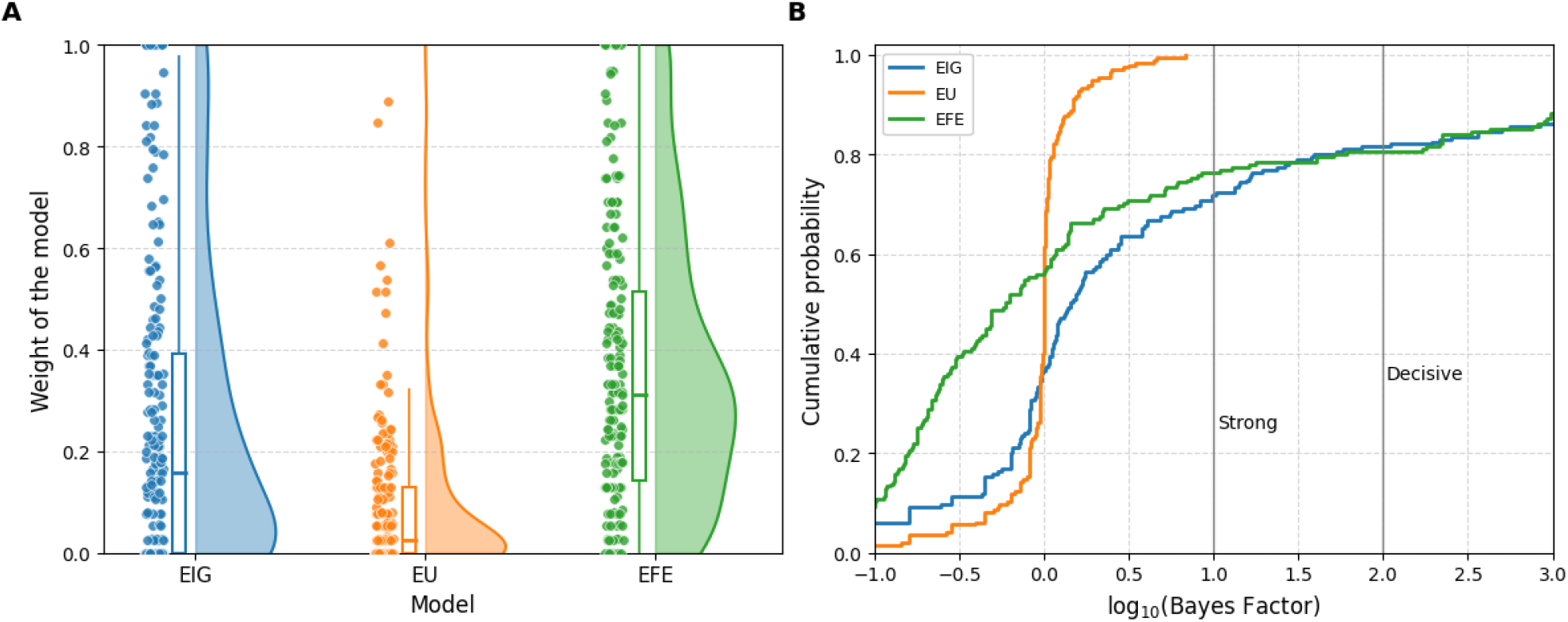
Random versus value-based choices. **(A)** Distribution of mixing weight *w* quantifying, for each individual and session, how much of participant’s behaviour is governed by structured value signals relative to random exploration: *w = 0* corresponds to pure random behaviour while *w = 1* corresponds to fully deterministic, value-driven choices. **(B)** Cumulative distribution of Bayes Factor (log10); the intercept with a BF of 1 and 2 indicates the proportion of sessions with either strong or decisive evidence in favour of EIG and EFE-based behavioural strategies, respectively.

To quantify such effects, we computed the Bayes Factor (based log10) comparing a model with random exploration with three models including the EIG, EU and EFE valuation systems. In the current analyses, the Bayes Factor (BF) indicates the strength of evidence in favor of each model (EIG, EU and EFE) with respect to purely random agents. A value from 1 to 2 indicates strong evidence, and larger than 2 a decisive evidence. Figure 3B shows the cumulative distribution of BFs. It shows that no sessions can be associated with strong evidence in favor of a EU-based model (orange curve). On the other hand, approximately 30% of sessions display a strong contribution of EIG and EFE (blue and green curves, respectively) and approximately 20% of sessions provide decisive evidence.

### Brain areas encoding expected information gain

The behavioural and computational modelling results suggest that EIG is a relevant metric to account for participants’ behavioral policies in information seeking in our task. We then investigated whether EIG is present in the brain and which areas it recruits. To address this question, we performed information-theoretical analyses of source-level high-gamma activity (HGA, from 60 to 120Hz) from magnetoencephalography (MEG). We focused on broadband gamma for two main reasons. First, it has been shown that the gamma band activity correlates with both spiking activity and the BOLD fMRI signals (Mukamel et al. 2005; Niessing et al. 2005; Lachaux et al. 2007; Nir et al. 2007; Ray and Maunsell 2011). Moreover, it is commonly used in MEG and SEEG studies to map task-related brain regions (Brovelli et al. 2005; Crone et al. 2006; Jerbi et al. 2009). Therefore, the analysis of HGA facilitates making a bridge with complementary techniques such as neuroimaging (fMRI) and invasive neurophysiological data from non-human primates. Second, single-trial and time-resolved HGA can be exploited for the analysis of cortico-cortical interactions in humans using MEG and iEEG techniques (Brovelli et al. 2015, 2017; Combrisson, Allegra, et al. 2022; Combrisson et al. 2024, 2025), thus providing an efficient marker for brain connectivity analyses. To characterise both the spatial organisation and temporal dynamics of brain areas encoding learning signals, we performed time-resolved mutual information analyses between the across-trials high-gamma activity (HGA) and expected information gain (EIG). Group-level random-effect analyses, combining permutation tests and cluster-based statistics (see Methods), were used to identify temporal clusters displaying significant encoding of learning signals and HGA. We observed that HGA in the left sensorimotor regions, including the motor and dorsal premotor regions, and the somatosensory cortex contained information about EIG at the time of action execution (left panel in Fig. 4). More specifically, the precentral gyrus and sulcus associated with the upper limb (dorsal portions) showed the strongest correlation with EIG. The regions correspond to the dorsal premotor and primary motor cortices. In addition, the postcentral gyrus and sulcus have significant activation, corresponding to the primary somatosensory cortex of the lateral parietal lobe (right panel in Fig. 4). Both activations were exclusively observed in the left hemisphere, consistent with the motor response with the right hand. These results suggest that the motor system carries information about EIG around action in the motor system for motor selection and execution, and in the somatosensory system for the perception of touch and movement. This indicates that the brain evaluates how much it would learn from each action within the very circuits that prepare those actions, and that information seeking is an embodied computation, not just a cognitive one.

**Figure 4.**
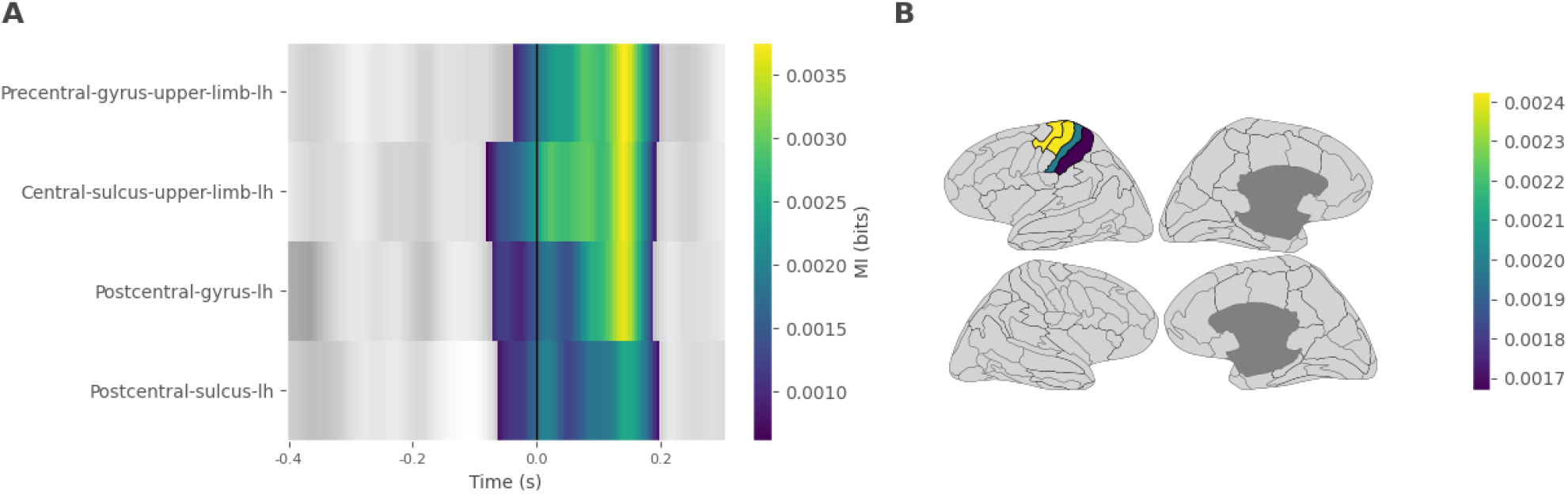
Neural correlates of expected information gain (EIG) for the chosen action. Four brain regions showed significant encoding of EIG around the time of motor response (panel **A**) and spatially clustered in the sensorimotor regions of the left hemisphere (panel **B**).

### Cortico-cortical broadcasting of expected information gain

We then investigated where information about EIG is generated and where it propagates. To do so, we used a recently-developed information-theoretic measure termed Feature-specific Information Transfer (FIT). Feature-specific Information Transfer (FIT) quantifies how much information about specific features flows between brain areas (Celotto et al., 2023). FIT merges the Wiener-Granger causality principle (Granger, 1980; Brovelli et al., 2004; Bressler and Seth, 2011) with content specificity based on the PID framework (Williams and Beer, 2010; Lizier et al., 2018). FIT isolates information about a specific task variable Y (expected information gain) carried by the current activity of a receiving neural population, which was not contained in its past activity, and which was instead present by the past activity of the sender neural population. We used the FIT measure to quantify the broadcasting of EIG between the four brain regions displaying local encoding. Two directional relations significantly contained information about EIG and broadcast information from the motor to the somatosensory regions (Fig. 5). The temporal information is aligned from the point of view of the receiving brain area. The significant EIG-specific information transfer lasted approximately 0.1s around action execution. The main source of information about EIG was the dorsal premotor and motor region in the precentral gyrus, and it broadcasted information about EIG to both the somatosensory areas in the postcentral gyrus and sulcus (Fig. 5).

**Figure 5.**
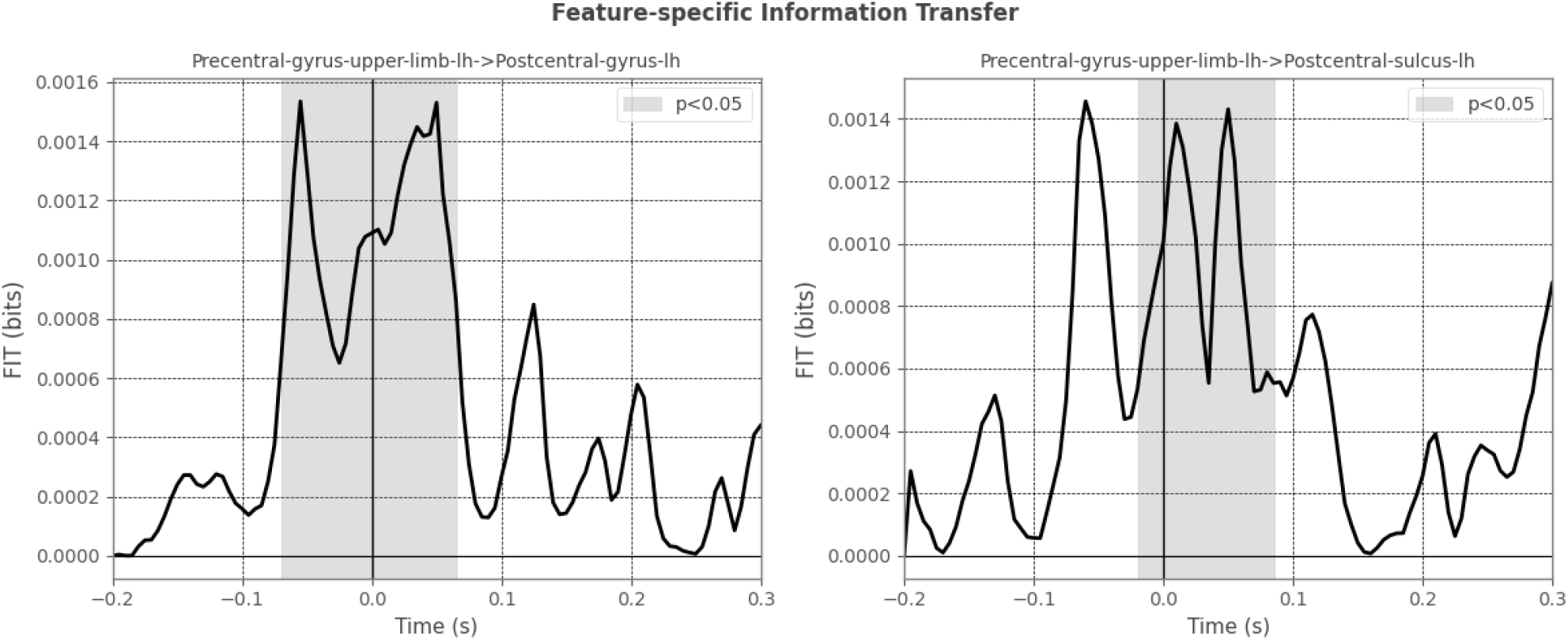
Broadcasting of expected information gain (EIG) in the brain. Two cortico-cortical interactions significantly carried information EIG. The broadcasting of information brain regions showed significant encoding of EIG around the time of motor response (left panel) and spatially clustered in the sensorimotor regions of the left hemisphere (right panel).

## DISCUSSION

### Dissociating information-seeking from reward maximisation during goal-directed causal learning

In the current study, we examined the ability to learn beliefs about action-outcome relationships, which constitutes a foundation for adaptive agency, rational decision-making, and flexible behavioral adjustment across changing environments and social contexts (Bandura, 1997). The acquisition of action-outcome causal representations is supported by the goal-directed system (Tolman, 1948; Dickinson, 1994; Dolan & Dayan, 2013). Although progress has been made towards the characterisation of underlying reward and information maximisation processes (Cohen et al. 2007; Schwartenbeck et al. 2019; Gottlieb et al. 2013; Cockburn et al. 2022; Badre et al. 2012; Frank et al. 2009; Mehlhorn et al. 2015), dissociating these learning objectives is not trivial. Indeed, they typically co-occur in both naturalistic and laboratory settings shaped by reinforcement schedules and bandit tasks, in which participants maximize cumulative reward by sequentially sampling from options with different payoff distributions (Daw et al. 2006; Cohen et al. 2007). A central contribution of this work is the design of a task that isolates the acquisition of causal beliefs from the pursuit of extrinsic rewards during goal-directed learning. The volleyball paradigm introduced here (Fig. 1A), circumvents this confound by instructing participants to evaluate the causal impact of a new player on team performance across simulated matches without incentivising match wins. In this task, participants sampled both Play and Not-Play actions to estimate the conditional probabilities needed to compute the action-outcome contingency ΔP (Jenkins and Ward 1965; Allan 1980; Allan and Jenkins 1980; Morris et al. 2017). The observed near-uniform probability of choosing Play across trials and contingency levels (approximately 0.5, Fig. 2C) confirms that participants adopted an exploratory strategy consistent, and is incompatible with reward-maximising policies that would strongly favour the action associated with the highest win probability. In addition, the significant correspondence between participants’ causal judgements and the contingency ΔP predicted by the model (R^2^ = 0.85, Pearson r = 0.92, p < 0.001, Fig. 2B) suggests near-optimal trial-by-trial belief updating consistent with an ideal Bayesian observer (Fig. 2A). Overall, these results provide evidence that our tasks disentangles information-seeking from reward maximisation during goal-directed causal learning.

### Expected information gain as a computational objective of action selection for explorative behaviors during for goal-directed causal learning

We then addressed an open question concerning the underlying computational objectives that shape the behavioral strategies participants use to learn causal action–outcome relationships. Current models from reinforcement learning and Bayesian learning (Friston et al. 2015, 2017; Sutton and Barto 2018) converge on the notion that sampling informative options is a core computational objective in goal-directed causal learning. In humans, explorative information sampling has been shown to rely on a mixture of random and directed strategies (Wilson et al. 2014; Gershman 2018). Random exploration, characterised by stochastic action selection independent of expected value, is thought to reflect “injected” noise in action selection policies and it has been argued to facilitate sampling in environments with structured uncertainty. Directed exploration, by contrast, reflects the assignment of an uncertainty bonus to under-sampled options and has been formalised within the Active Inference framework as the maximisation of expected information gain (Friston et al. 2015; Parr et al. 2022). This notion also aligns with information maximisation accounts of novelty seeking and exploratory behaviour (Kidd and Hayden 2015; Gottlieb et al. 2013; Gottlieb and Oudeyer 2018; Schwartenbeck et al. 2019; Modirshanechi et al. 2025).

To quantify the contribution of random and directed exploration, we modelled the probability of performing an action as a weighted mixture of two components: a random baseline and a general value-driven component. In other words, we modelled participants’ choices as resulting from a tradeoff between randomly sampling actions and a directed value-based component. We tested predictions from Active Inference (AI), in the context of goal-directed learning and planning, which suggests that agents select actions to minimize the expected free energy of future outcomes, which is equal to the sum expected utility (extrinsic value) and expected information gain (intrinsic value) (Friston et al. 2015). For both the EIG, expected utility (EU), and expected free energy (EFE) models, the distribution of mixing weights *w*, quantifying how much of each participant’s behaviour is governed by value signals relative to random exploration, was skewed towards zero (Fig. 3), indicating that choices were dominated by random behaviour. However, a substantial proportion of sessions showed weights equal to or exceeding 0.5 for EIG and EFE, indicating that directed and epistemic exploration contributed meaningfully to behaviour in these cases (Fig. 3A, blue and green), which approximately 20% showing decisive evidence (log_10_ BF > 2) in favour of EIG and EFE over a purely random agent, while no sessions provided equivalent evidence for a EU-based policy (Fig. 3B). The near-identical BF distributions for EIG and EFE was expected given that EFE reduces to EIG when EU is negligible, which is a direct consequence of the Volleyball task design.

Overall, the results suggest that EIG is a computational objective for action selection during explorative behaviors in goal-directed causal learning, and it provides support to previous modelling frameworks formalising information seeking and exploration using model-based reinforcement algorithms or Bayesian models (Gläscher et al. 2010; Liakoni et al. 2021; Faraji et al. 2018; Yu and Dayan 2005; Modirshanechi et al. 2022) and epistemic value components predicted by Active Inference (Friston et al. 2015; Parr et al. 2022).

### Sensorimotor encoding of expected information gain

The next contribution of our study has been the investigation of the neural bases of expected information gain (EIG). To do so, we used human magnetoencephalography (MEG) and information-theoretical analyses of source-level high-gamma activity (HGA). The results show that expected information gain (EIG) is not only a computational objective of exploratory behavior, but is also significantly contained in the brain at the time of action execution (Fig. 4A). The brain regions showing significant correlation with EIG surprisingly confined to a single cluster of adjacent cortical areas. These included the precentral gyrus and central sulcus, corresponding to the dorsal premotor and primary motor cortices, as well as in the postcentral gyrus and sulcus, corresponding to the primary somatosensory cortex. All activations were confined to the left hemisphere, contralateral to the right hand used for action execution (Fig. 4B).

These results further support the view that action selection during goal-directed causal learning is partly governed by the epistemic value of actions (Friston et al. 2015). Accordingly, agents select actions with the highest expected information gain in order to reduce uncertainty about the causes of valuable outcomes. The results suggest that the neural substrate for active sampling based on EIG computations resides in the sensorimotor regions.

This spatial pattern contrasts with well-established cortical circuits for computing and comparing expected utility during action selection, including the orbitofrontal cortex, ventromedial prefrontal cortex, and striatum (Padoa-Schioppa and Assad 2006; Ballesta et al. 2020; Gardner et al. 2020; Gore et al. 2023; Daw and O’Doherty 2014; Rangel et al. 2008). The sensorimotor encoding of EIG suggests a fundamentally different organisation for epistemic valuation, particularly when it is directly linked to motor action selection. We suggest that the sensorimotor cortex serves as the computational site where the epistemic value of forthcoming actions is evaluated, because it is the system that will implement those actions. According to predictive processing accounts of motor control, the sensorimotor cortices continuously generate predictions about the sensory consequences of actions, rather than motor commands (Adams et al. 2013). Agents create a probabilistic model of how sensory inputs are caused by actions (e.g., the winning or losing icons in the volleyball task), such that the resulting predictions can guide active sampling of sensory data. Our results support a sensorimotor encoding of EIG for active sampling of actions, where the sensorimotor system simulates action outcomes to evaluate their informativeness. The computation of EIG in the sensorimotor system is a natural extension of the forward modelling computations already performed by sensorimotor and premotor circuits.

Previous results have proposed that reward and information value recruit a common neural circuitry in the limbic system of non-human primates (Kobayashi and Kable 2024; Levy and Glimcher 2012), OFC in human fMRI (Charpentier et al. 2018) and in frontal EEG electrodes (Brydevall et al. 2018). These areas are main targets of dopaminergic projections, which contain expected advance information about future outcomes in non-instrumental settings (Bromberg-Martin and Hikosaka 2009; Bromberg-Martin et al. 2024; Bromberg-Martin and Hikosaka 2011). The anterior cingulate cortex (ACC) and two subregions of the basal ganglia (the internal-capsule-bordering portion of the dorsal striatum and the anterior pallidum), signalling reward uncertainty and information-anticipatory activity (White et al. 2019), may additionally provide a substrate for common encoding of reward and information value. Our results challenge the view that information seeking exclusively relies on the neural circuitry for reward-based learning, and it provides evidence for a crucial role of the motor system, at least in goal-directed causal learning.

### Cortico-cortical broadcasting of epistemic value

In order to investigate how information about EIG propagates in the sensorimotor system, we performed directional information-theoretic analyses using a recently-developed metric termed Feature-specific Information Transfer (FIT). FIT quantifies how much information about task-related or cognitive variables flows between brain areas (Celotto et al., 2023), by merging the Wiener-Granger causality principle (Granger, 1980; Brovelli et al., 2004; Bressler and Seth, 2011) with content specificity based on the PID framework (Williams and Beer, 2010; Lizier et al., 2018). The results showed that information about EIG propagates from the dorsal premotor and primary motor cortex in the precentral gyrus to both the postcentral gyrus and the postcentral sulcus around the time of action execution. This observation suggests that the motor system, once computed the epistemic value of an intended action and committed to a motor plan, send this information to somatosensory cortex to the anticipation not merely the tactile and proprioceptive consequences of the movement, but also its epistemic consequence (i.e., upcoming actions outcome). This process may implement a form of epistemic efference copy embedded within the sensorimotor hierarchy, and allow subsequent updating. The directionality of this transfer and the absence of significant EIG broadcast in the reverse direction support a hierarchical organisation consistent with predictive cording hypothesis that higher levels of the sensorimotor hierarchy generate predictions that are passed to lower levels for implementation and error correction (Adams et al. 2013; Friston et al. 2010). Within such a framework, the motor system, which is agranular (that is, it lacks a fully expressed granular layer IV), is thought to issue motor predictions to the spinal cord as deterministic commands to move the body. Simultaneously, the motor system is thought to send somatosensory predictions to the primary somatosensory cortex to represent the anticipated proprioceptive and kinaesthetic consequences of the movement. These somatosensory predictions are akin to ‘efference copies’ of corollary discharge that enable the brain to track self-initiated movements (Shipp et al. 2013). We suggest that the sensorimotor systems additionally participates by predicting the sensory causal effect of actions (sensory states, such as the winning or losing stimuli in the volleyball task) during goal-directed causal learning.

### Relation to first-personal causal understanding and embodied cognition

The goal-directed learning task used in the current study assesses a cognitive ability at the interface between reinforcement and contingency learning. During the development of human causal learning and reasoning, the earliest signs of such ability are present at infancy (such as “crying to attract attention” in newborns) and they are shared with some non-human animals (“Expecting that if I shake the branches, then I will make the fruit fall”). These forms of causal understanding are referred to as first-personal and they are thought to be tied to the agent’s own goal-directed action system (Goddu and Gopnik 2024). The convergence of localized encoding and directed functional connectivity within the sensorimotor system is consistent with a first-person, action-centered account of causal learning. In addition, these findings challenge the canonical view that an abstract computational signal such as expected information gain (EIG), which presupposes integration of probabilistic beliefs about action–outcome contingencies and prospective information states, is exclusively represented within the reward-based goal-directed valuation network. Instead, they suggest that the intrinsic value or informational value of actions is not computed in a dedicated valuation module and then relayed to the motor system; rather, the motor system itself tracks what it would learn from acting, and this epistemic computation is made available to the somatosensory system via cortico-cortical broadcast. Under this view, evaluating the informativeness of an action is intrinsic to the act of representing and selecting among possible actions, rather than a process that occurs prior to or independently of motor planning. This interpretation represents a strong form of embodied epistemic valuation that, to our knowledge, has not previously been supported by empirical data in the context of goal-directed causal learning. Our findings are more consistent with embodied cognition accounts, which propose that cognitive processes are fundamentally constituted by the sensorimotor systems through which agents interact with their environments. Nevertheless, our results may not be generalisable to different contexts, such as learning by observing others’ goal-directed actions (third-personal causal learning), or even by generalizing from the consequences of one’s own accidental movements.

### Open questions about information valuation system and complementary learning rule

The finding that EIG is represented in the sensorimotor system has important theoretical implications. It complements the dominant view that the brain’s valuation systems are primarily organised around extrinsic reward signals encoded in the orbitofrontal cortex and ventromedial prefrontal cortex (Daw and O’Doherty 2014; Rangel et al. 2008; Padoa-Schioppa and Assad 2006; Rushworth et al. 2011). In addition, it suggests that the neural system dedicated to intrinsic valuation spans a broader brain network than previously thought, which was shown to include the ventral striatum, ventromedial prefrontal cortex (vmPFC), and dorsolateral prefrontal cortex (Bromberg-Martin and Hikosaka 2009; Bromberg-Martin et al. 2024; Bromberg-Martin and Hikosaka 2011; Kobayashi and Hsu 2019; Kobayashi et al. 2021; Kobayashi and Kable 2024). It supports recent findings that information valuation relies on distributed cortical circuits, including visual, parietal, lateral prefrontal, and ventromedial/orbitofrontal regions, which interact through higher-order functional dynamics and project to reward circuitry (Combrisson et al. 2025). A recent theoretical formulation has proposed that EIG is computed by the noradrenergic (NA) system (“OSF,” n.d.). Within this framework, the NA is proposed to act as a higher-level meta-controller that dynamically adjusts lower-level behavioral parameters, including exploration rate, the balance between goal-directed and habitual control, learning rate, and cognitive flexibility. It remains speculative, however, whether the diffuse projection of the locus coeruleus (LC) noradrenaline neurons to sensorimotor regions, including motor cortex and somatosensory cortex, plays a role in the neuromodulatory effect of epistemic value.

A second open question regards the relation to computational models proposing learning noise as a source of exploratory (non-greedy) decisions (Findling and Wyart 2024; Findling et al. 2019). Learning noise is defined as an unintentional, undirected quantity and random deviation from the exact application of the learning rule, scaling with prediction error magnitude. To our knowledge, the contribution of learning noise and epistemic value to exploratory behavior remains unclear, as do their underlying neural circuits.

### Conclusions

The present study demonstrates that expected information gain is a behaviourally relevant intrinsic valuation signal during goal-directed causal learning, and that this signal is contained and broadcast within the sensorimotor system rather than exclusively within canonical prefrontal valuation circuits. The combination of Bayesian computational modelling, capturing trial-by-trial belief updating via an ideal Bayesian observer, with information-theoretic analyses of MEG high-gamma activity and Feature-specific Information Transfer (Celotto et al. 2023) provides a multi-level account of how epistemic value is computed and communicated in the human brain. These results support a view of epistemic valuation as an embodied process, in which the evaluation of the informational consequences of actions is performed within the very neural circuits that represent and prepare those actions (Friston et al. 2015; Parr et al. 2022). By grounding cognitive computations about causal uncertainty in sensorimotor systems, the brain may achieve an efficient integration of epistemic and motor planning processes that supports flexible, curiosity-driven exploration of causal environments (Gottlieb and Oudeyer 2018; Schwartenbeck et al. 2019). These findings have implications not only for the neuroscience of causal learning and active inference, but also for computational models of artificial agents that aspire to implement human-like causal reasoning through embodied action and world models (Hafner et al. 2025).

## MATERIALS AND METHODS

### Goal-directed causal learning task and experimental conditions

Thirteen healthy participants accepted to take part in our study, all of them were right handed, eight were females and five males, and the average age was around 25 years. All participants gave written informed consent according to established institutional guidelines, and they received monetary compensation (50 euros) for their participation. The project has been promoted by the INSB of the CNRS (CNRS No. 17020; ANSM No. 2017-A03614-49) and approved by the ethical committee CPP Sud-Méditerranée.

To dissociate reward maximization from information-seeking processes, we designed a task that required participants to learn the causal effect of their actions in different scenarios of varying contingencies, without incentivizing reward maximization, thereby isolating learning about action–outcome causality. The task was inspired by previous neuroimaging studies (Liljeholm et al. 2011). Participants were instructed to play a videogame in which they had to impersonate a trainer of a volleyball team. Their goal was to evaluate the causal effect of a new player in their team by means of a simulator. To do so, they could simulate a set of forty fictitious matches. For each match, the participant (impersonating the trainer of the team) could choose to either add the new player to the team or not. Each fictitious match represented a trial in the task. Each learning session started with the presentation of two cues associated with two options: one cue (depicting a “Play” symbol) was associated to the simulation of a match with the new player (“Play”), whereas the other cue (depicting a “Pause” symbol) was associated to a match without the new player “Not Play”). Participants chose using the right and left buttons, which were associated with either the “Play” or “Not Play” choices (Fig. 1A). The position of the “Play” and “Not Play” cues did not vary during a learning block, but it was randomly assigned across blocks to avoid possible biases due to positional effect. The outcome of the fictitious match was then presented 0.3s after the response of the participant. The delay duration was fixed, rather than random as often used in decision making tasks, to introduce a stronger causal relation between action and outcome. The outcome cue was either a green happy or a red sad face presented at the center of the screen, and informed the participants that the match either played or not played by the new player was won or lost, respectively. The outcome was displayed for 1.5 s. After outcome disappearance, participants could simulate another match. The task design was self-paced, meaning that no visual cue instructed participants to perform a choice.

During a learning session of forty matches, the causal effect of the new player in the team was dictated by the contingency Δ*P* defined as the difference between the probability of winning the match given the presence of the new player P(Win | Play) and the probability of winning given the absence of the player P(Win | Not Play). A positive Δ*P* is associated with a player with positive causal effect, whereas a negative Δ*P* is associated with a player with negative causal effect. At the end of each learning block, participants had to verbally report a causal score ranging from −100 to 100 associated with the new player. A causal score of −100 would correspond to players that make the team lose every time they play, whereas a score of 100 corresponds to players that would make the team win every time they played. A score of 0 would correspond to a player with null causal effect on the result of the match. A total of 15 learning blocks associated with 15 different new players was submitted to the participants. Participants were instructed that learning blocks were independent. Each new player was associated with a given Δ*P*. Five values of Δ*P*were tested (−0.6, −0.3, 0, 0.3 and 0.6). Each of the five levels of Δ*P* was composed of three combinations of conditional probabilities (Fig. 1B), thus giving a total of fifteen learning blocks. Participants performed fifteen learning blocks in a randomized order, and all of them received the same instructions.

### Computational model of goal-directed causal learning

#### Bayesian ideal observer

We assume that the first element of goal-directed causal learning is the creation and update of internal beliefs of action-outcome conditional probabilities and contingency values during learning. We modeled the inference of causal action–outcome relations within a Bayesian inference framework. Bayesian inference is a principled statistical model based on Bayesian updating of beliefs about the current state of the world. In our task, the causal effect of a new player is quantified as the contingency:

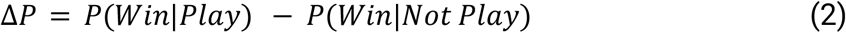

We should note that ΔP is equivalent to the Average Causal Effect (ACE) as defined in Pearl’s interventional framework and do-calculus: ACE = P(Y = 1|do(X = 1)) − P(Y = 1|do(X = 0)). This equivalence holds under the assumption that the experimental conditions approximate ideal interventions, thus supporting the use of this framework for the study of causal learning. Each conditional probability is estimated separately. Since match outcomes *y*_*t*_ across trials *t* are binary random variables (win and loss), observations are governed by the Bernoulli distribution *p*(*y*_*t*_ |θ_*t*_) = *Bernoulli*(θ_*t*_). Prior beliefs about the probability of winning θ_*t*_ are represented by a Beta distribution *p*(θ_*t*_ |*a*_*t*_) = *Beta*(α_*t*−1_ (*a*_*t*_), β_*t*−1_ (*a*_*t*_)), where *a*_*t*_ denotes the selected action (Play and Not-play) at trial *t*. The shape hyperparameters α_*t*_ and β_*t*_ are action dependent and control the expectation and precision of prior beliefs about θ_*t*_. We considered α_0_ = β_0_ = 1. 1 for every action, producing a symmetric prior centered at 0.5 with mild regularization. Because the Beta distribution is conjugate to the Bernoulli distribution, combining the prior with the likelihood via Bayes’ theorem yields a posterior that remains in the Beta family, with parameters updated by adding observed successes and failures. The posterior, after observing *y*_*t*_, is given by:

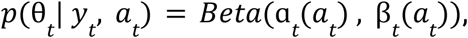

where

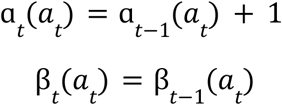

if *y*_*t*_ is a win (success) when choosing *a*_*t*_, and

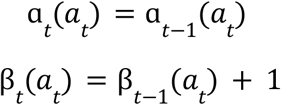

if *y*_*t*_ is a loss (failure) when choosing *a*_*t*_.

Given the properties of the Beta distribution, at each trial, the mean estimate of θ_*t*_ is:

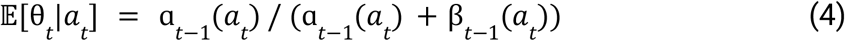

The contingency estimate was computed as the difference between posterior means. This formulation provides a parsimonious and analytically coherent account of Bayesian learning from binary outcomes.

#### Action valuation mechanisms

The second element of goal-directed causal learning is the selection of actions based on intrinsic or extrinsic values. Here, we tested notions from Active Inference (AI) as a leading model of intrinsic reward valuation. Active Inference is a normative model of perception, action and planning in terms of variational free energy minimization (Parr et al. 2022). In the context of goal-directed learning and planning, AI predicts that agents select actions to minimize the expected free energy of future outcomes. Since the negative free energy (or quality of a policy) can be decomposed into extrinsic and epistemic (or intrinsic) value, minimizing expected free energy is equivalent to maximizing extrinsic value (or expected utility, defined in terms of prior preferences or goals), while maximizing information gain or intrinsic value (or reducing uncertainty about the causes of valuable outcomes or epistemic value) (Friston et al. 2015). In the current causal learning task, Active Inference predicts that the intrinsic value of each action (Play and Not-Play) is proportional to the Expected Information Gain (EIG), which reflects the epistemic value. In Bayesian statistics, information gain can be defined as the amount of information required to “revise” one’s beliefs from the prior probability distribution to the posterior probability distribution. This measure is formalised via the Kullback–Leibler divergence *D*_*KL*_ between the posterior and prior distributions. In neuroscience, this measure is also referred to as the Bayesian surprise (Itti and Baldi 2009; Modirshanechi et al. 2023). The EIG is therefore defined as the mean information gain averaged over possible future outcomes when selecting action *a*_*t*_, as:

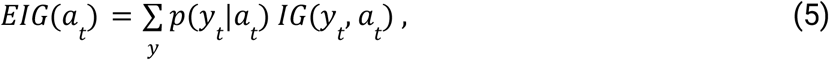

where the information gain for a specific pair of action-outcome is:

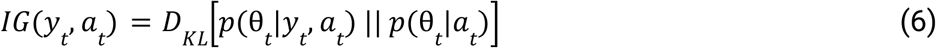

The agent is assumed to choose the action associated with the highest EIG.

Although the learning task was designed to specifically promote information seeking and action selection based on expected information maximisation, we compare the EIG model with two complementary models based on standard subjective Expected Utility (EU) and Expected Free Energy (EFE). The Expected Utility (EU) of an action is defined as the probability-weighted sum of the utility of outcomes:

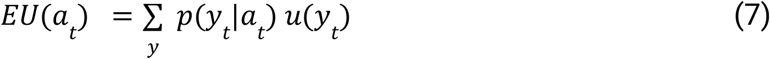

We adopt the utility function used in Active Inference, where the utility of outcomes is defined as the logarithm of their softmax-normalized rewards *c*(*y*_*t*_), given by:

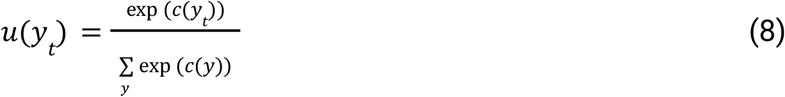

where *c*(*win*) = 1 and *c*(*loss*) =− 1. The Expected Free Energy (EFE) as predicted by Active Inference is the defined as:

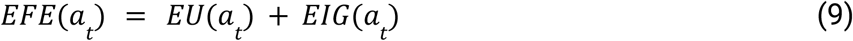

#### Action selection mechanisms

The third component of the model is the action selection process based on acquired heuristics and/or valuation system. In the setting of the Volleyball task, where the two unknown conditional probability distributions must be estimated under a constrained sampling limit (40 trials), and under symmetric priors, the expected posterior risk (e.g., mean squared error) remains minimized by approximately equal allocation of choices across the two alternatives. In order to quantify the contribution of the abovementioned valuation processes, we took a baseline model, an agent that samples each action randomly with no learning and no valuation systems. We modelled the probability of performing an action as a weighted mixture of two components: a random baseline and a value-driven component. Formally, at each trial the probability of *a*_*t*_ is given by

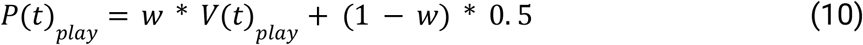

where *w* ∈ [0, 1] is the weight assigned to the value-driven component and (1−*w*) is the weight assigned to pure random behaviour (Bernoulli with *p* = 0.5). The value-driven component *V*_*play*_ is derived from an argmax policy applied to a model-specific value signal, which can reflect the expected reduction in uncertainty about the environment regardless of reward (EIG model), the expected reward obtained by playing (EU model) or a combination of both (EFE model). The mixing weight *w* thus quantifies, for each individual, how much of their behaviour is governed by structured value signals relative to random exploration: *w* = 0 corresponds to pure random behaviour while *w* = 1 corresponds to fully deterministic, value-driven choices.

### Model fitting and model comparison

The parameterisation in Eq. 10 ensures that the random model is nested within every model of learning-related valuation, as the special case w = 0, enabling fair model comparison via BIC. Each model’s free parameters were estimated by maximum likelihood (MLE) applied to the sequence of observed actions. The free parameters are w and the reward preference weight *C*. For the EU model, the reward preference weight *C* can be set fixed, because the model’s decisions with the argmax are identical for any choice of *C*. Thus, for the one-parameter models (EIG, EU), MLE over *w* was performed using the bounded Brent method (scipy.optimize.minimize_scalar). This is exact because, for fixed argmax predictions, the log-likelihood LL(*w*) = ∑_t_ log[*w* · P_argmax,t_ + (1−*w*) · 0.5] is strictly concave in *w*, guaranteeing convergence to the global optimum. For the two-parameter EFE model (*w, C*), the objective function is piecewise constant in *C*, because ΔEFE_t_(*C*) = 2 Δ*p*_*t*_^W^ · *C* + ΔEIG_*t*_ is affine in *C*; thus, the argmax pattern changes only at breakpoints *C*_*t*_^*^ = −ΔEIG_t_ /(2 Δ*p*_*t*_^W^). The global MLE was therefore found by an analytical breakpoint search: all breakpoints within the parameter range *C* ∈ [10^−3^, 10] were enumerated, one representative *C* value was selected per constant-argmax interval, and *w* was optimised exactly via Brent’s method for each candidate *C*; the pair (*w, C*) yielding the highest log-likelihood was retained.

Model comparison was performed by computing the Bayesian Information Criterion (BIC = *k* log *n* − 2 LL, where *k* is the number of free parameters and *n* the number of trials), which penalises model complexity and permits fair comparison across models with different numbers of parameters. Then, to compute the strength of evidence in favor of each model, we computed the log_10_ Bayes factor between each model and the random agent, which can be approximated from their BIC values as

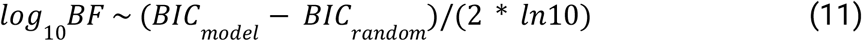

where a value from 0.5 to 1 indicates substantial evidence, a value from 1 to 2 a strong evidence, and larger than 2 a decisive evidence.

### MEG and MRI recordings, cortical parcellation and source model

Magnetoencephalographic (MEG) recordings were performed using a 248 magnetometers system (4D Neuroimaging magnes 3600). Visual stimuli were projected using a video projection and motor responses were acquired using a LUMItouch® optical response keypad with five keys. Presentation® software was used for stimulus delivery and experimental control during MEG acquisition.

Anatomical T1-weighted MRI images were acquired for all participants using a 3-T whole-body imager equipped with a circular polarised head coil (Bruker). Single-subject cortical parcellation was performed using the Virtual Epileptic Patient (VEP) atlas (Wang et al. 2021). After denoising using non-local means (Coupe et al. 2008), T1-weighted MR-images were segmented using the FreeSurfer “recon-all” pipeline (v5.3.0). The VEP atlas is a subdivision of the Destrieux parcellation of the cortex (Destrieux et al. 2010), which gives a cortical subdivision based on major gyri and sulci. The creation of the VEP atlas is automated and it produces 146 cortical parcels. Splitting and merging operations are performed using the vertices of the triangulated cortical surface and then projected to the voxels using FreeSurfer’s mri_aparc2aseg tool. Such parcellation has a spatial resolution adequate for both intracranial SEEG (Wang et al. 2021) and MEG studies (https://github.com/HuifangWang/VEP_atlas_shared).

Finally, we created source space and forward models readable by MNE-python (Gramfort et al. 2013) using cortical meshes generated using VEP atlas and generated by Freesurfer. These elements are needed for the power estimation at the source level, which will be discussed in the next paragraph. The anatomical meshes were coregistred with the MEG data using the ‘mne coreg’ interface.

### Single-trial and atlas-based high-gamma activity (HGA)

We focused on broadband gamma from 60 to 120Hz for two main reasons. First, it has been shown that the gamma band activity correlates with both spiking activity and the BOLD fMRI signals (Mukamel et al. 2005; Niessing et al. 2005; Lachaux et al. 2007; Nir et al. 2007; Ray and Maunsell 2011), and it is commonly used in MEG and iEEG studies to map task-related brain regions (Brovelli et al. 2005; Crone et al. 2006; Jerbi et al. 2009). Therefore, focusing on the gamma band facilitates linking our results with the fMRI and spiking literature on probabilistic learning. Second, single-trial and time-resolved high-gamma activity can be exploited for the analysis of cortico-cortical interactions in humans using MEG and iEEG techniques (Brovelli et al. 2015, 2017; Combrisson, Allegra, et al. 2022; Combrisson et al. 2024, 2025).

MEG signals were high pass filtered to 1Hz, low-pass filtered to 250 Hz, notch filtered in multiples of 50Hz and segmented into epochs aligned on stimulus and outcome presentation. Independent Component Analysis (ICA) was performed to detect and reject cardiac, eye-blink and oculomotor artefacts. Artefact rejection was performed semi automatically, at first by a visual inspection of the epochs’ time series, then by means of the *autoreject* python library (Jas et al. 2017) to detect and reject bad epochs from further analysis.

Spectral density estimation was performed using a multi-taper method based on discrete prolate spheroidal (slepian) sequences (Percival and Walden 1993). To extract high-gamma activity from 60 to 120Hz, MEG time series were multiplied by *k* orthogonal tapers (*k* = 11) (0.2s in duration and 60Hz of frequency resolution, each stepped every 0.005s), centred at 90Hz and Fourier-transformed. Complex-valued estimates of spectral measures 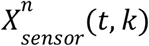, including cross-spectral density matrices, were computed at the sensor level for each trial *n*, time *t* and taper *k*. Source analysis requires a physical forward model or leadfield, which describes the electromagnetic relation between sources and MEG sensors. The leadfield combines the geometrical relation of sources (dipoles) and sensors with a model of the conductive medium (i.e., the head model). For each participant, we generated a boundary element model using a single-shell model constructed from the segmentation of the cortical tissue obtained from individual MRI scans (Nolte, 2003). Lead-fields were not normalised. The head model, source locations and the information about MEG sensor position for both models were combined to derive single-participant leadfields. The orientation of cortical sources was set perpendicular to the cortical surface.

We used adaptive linear spatial filtering to estimate the power at the source level. In particular, we employed the Dynamical Imaging of Coherent Sources (DICS) method, a beamforming algorithm for the tomographic mapping in the frequency domain (Gross et al. 2001), which is a well suited for the study of neural oscillatory responses based on single-trial source estimates of band-limited MEG signals. At each source location, DICS employs a spatial filter that passes activity from this location with unit gain while maximally suppressing any other activity. The spatial filters were computed on all trials for each time point and session, and then applied to single-trial MEG data.

Single-trial power estimates aligned with outcome and stimulus onset were log-transformed to make the data approximate Gaussian and low-pass filtered at 50Hz to reduce noise. Single-trial mean power and standard deviation in a time window from −0.5 and −0.1 s prior to stimulus onset was computed for each source and trial, and used to z-transform single-trial movement-locked power time courses. Similarly, single-trial outcome-locked power time courses were log-transformed and z-scored with respect to baseline period, to produce HGAs for the prestimulus period from −1.6 to −0.1 s with respect to stimulation for subsequent functional connectivity analysis. Finally, single-trial HGA for each brain region of the VEP atlas was computed as the mean z-transformed power values averaged across all sources within the same region. Single-trial HGA estimates were performed using home-made scripts based on MNE-python (Gramfort et al. 2013).

### Cortical areas encoding expected information gain

We used information-theoretic metrics to quantify the statistical dependency between single-trial HGA and expected information gain. Information-based measures quantify how much the neural activity of a single brain region explains a variable of the task. To this end, we computed the mutual information (MI); as a reminder, mutual information is defined as:

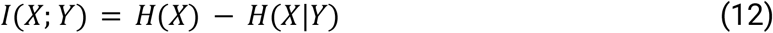

In this equation the variables *X* and *Y* represent the HGA power and the internal learning variables, respectively. *X* represents the HGA power over time, trials and regions. *Y* represents the evolution of EIG across trials. *H*(*X*) is the entropy of *X*, and *H*(*X*|*Y*) is the conditional entropy of *X* given *Y*. We used Gaussian-Copula Mutual Information (GCMI) (Ince et al. 2017), which is a semi-parametric binning-free technique to calculate MI. In the GCMI approach, which includes a rank based normalisation of the data, entropy values are computed using the differential entropy formula for a Gaussian distribution. The transformed data allows entropy estimation based on the determinant of the covariance matrix and the assumption of Gaussian marginals. The GCMI is a robust rank-based approach that allows detecting any type of monotonic relation between variables. More precisely, the GCMI is a lower-bound approximate estimator of mutual information for continuous signals based on a Gaussian copula, and it is of practical importance for the analysis of short and potentially multivariate neural signals. Note, however, that the GCMI does not detect non-monotonic (e.g., parabolic) relations. In the current work, the GCMI was computed across trials between time-resolved HGA and outcome-related learning signals. The parametric GCMI estimation was bias-corrected using an analytic correction to compensate for the bias due to the estimation of the covariance matrix from limited data (i.e. here, limited number of repetitions or trials) (Ince et al., 2017). Since this parametric correction only depends on the number of trials, the same value is going to be used for both permuted and non-permuted data. Therefore, this bias correction only impacts the estimated effect size but has no effects on statistical results.

### Broadcasting of expected information gain in the brain

In order to investigate whether information about EIG is broadcasted across different cortical regions, we used a recently-developed information-theoretic measure termed Feature-specific Information Transfer (FIT) that quantifies how much information about specific features flows between brain areas (Celotto et al. 2023). FIT merges the Wiener-Granger causality principle (Granger 1980; Brovelli et al. 2004; Bressler and Seth 2011) with content specificity based on the PID framework (Williams and Beer 2010; Lizier et al. 2018). Briefly, the PID framework allows the decomposition of multivariate mutual information between a system of predictors and a target variable, and to quantify the information that several sources (or predictors) variables provide *uniquely, redundantly* or *synergistically* about a target variable. In the current study, we considered the case where the source variables are the across-trials HGA of two brain regions (X_1_ and X_2_) and the target variable is the trial-by-trial evolution of the learning variable (Y). If pairs of brain regions X_1_ and X_2_ carry the same information about learning variable Y, we refer to it as *redundant* information. If only one area X_1_ or X_2_ carries information about Y, it is referred to as *unique* information carried by X_1_ or X_2_ about Y, and vice-versa for Y. Finally, if neither of the variables alone provide information about Y, but they need to be observed together, we refer to it as *synergistic* information. In other words, the knowledge of any of the predictors separately does not provide any information about the target variable C. Analytically, the PID proposes to decompose the total mutual information between a pair of source variables X and Y and a target variable Z *I*(*X*_1_, *X*_2_; *Y*) into four non-negative components:

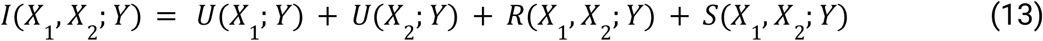

*U*(*X*_1_; *Y*) and *U*(*X*_2_; *Y*) are unique information carried for the two areas, respectively. *R*(*X*_1_, *X*_2_; *Y*) and *S*(*X*_1_, *X*_2_; *Y*) are the redundancy and synergy terms, respectively. In addition, the PID formulation links to classical Shannon measures of mutual information as

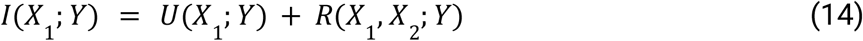

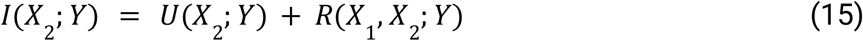

The problem with this approach is that its governing equations form an under-determined system: only three quantities that can be computed from the data (i.e., the mutual information quantities *I*(*X*_1_, *X*_2_; *Y*), *I*(*X*_1_; *Y*) and *I*(*X*_2_; *Y*)) for the four terms of the PID (i.e., two unique information, redundant and synergistic). To actually calculate the decomposition, an additional assumption must therefore be made. Here, we exploit the so-called minimum mutual information (MMI) PID, which has been shown to provide correct estimations for a broad class of systems following a multivariate Gaussian distribution (Barrett 2015; Luppi et al. 2022). According to the MMI PID, redundant information carried by pairs of brain regions is given by the minimum of the information provided by each individual source to the target,

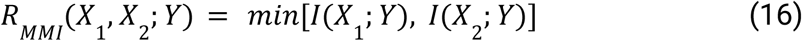

Then, synergistic information can be computed by substituting Eq. 14, 15, and 16 in Eq. 13 and rearranging the terms

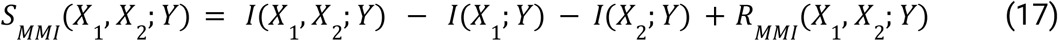

Equations 9 and 10 represent the redundant and synergistic information carried by the co-modulations in HGA of pairs of brain regions X_1_ and X_2_ about the learning variable Z, respectively. By definition, redundant functional interactions carry the same information about learning variables than the one carried by individual brain areas. On the other hand, synergistic connectivity reveals brain functional interactions that cannot be explained by individual contribution, but only by their combined and collective coordination.

Within the PID framework, FIT isolates information about a specific task variable Y in the current activity of a receiving neural population, which was not present in its past activity, and which was instead contained by the past activity of the sender neural population. More precisely, the FIT measure is based on two four-variables PID, and it is defined as the minimum of two “atoms”. The first atom is defined in a four-variable PID, which considers the learning variable *Y* as the target and three source variables, namely the brain activity in the past of the sending signal *X*_1_ (*t* − *τ*), the past of the receiver *X*_2_ (*t* − *τ*) and the present of the receiver *X*_2_(*t*). This atom formalises the principle of Granger-Wiener causality within the PID framework, and it adds the encoding of task variables (e.g., learning) in the time-lagged predictability between signals. More formally, the first atom is defined as the redundant information about *Y* carried by the past of *X*_1_ (*t* − *τ*) and the present of *X*_2_ (*t*), which is unique with respect to past of the *X*_2_ (*t* − *τ*).

We used the MMI PID to compute this atom, which was computed as

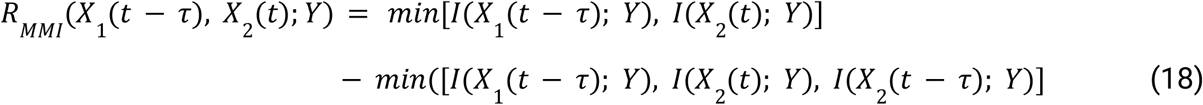

Note that the second term on the right-hand side assures the uniqueness with respect to the past of the receiver *X*_2_ (*t* − *τ*). The second atom for the calculation of the FIT is defined for a four-variable PID, considering the present of the receiver *X*_2_ (*t*), as the target variable and the learning variable *Y, X*_1_ (*t* − *τ*) and *X*_2_ (*t* − *τ*) as source variables. This second term assures that the FIT measure does exceed: i) the overall propagation of information between signals, referred to as the Directed Information (Massey 1990) or Transfer Entropy (Schreiber 2000), and quantified by means of the conditional mutual information *I*(*X*_1_ (*t* − *τ*); *X*_2_ (*t*)|*X*_2_ (*t* − *τ*)); ii) the mutual information between the learning variable and the past of the sender, *I*(*Y*; *X*_1_ (*t* − *τ*)) and iii) the mutual information between the learning variable and the present of the receiver, *I*(*Y*; *X*_1_ (*t*)). This term is defined as the redundant information about the present of the receiver *X*_2_ (*t*) carried by *X*_1_ (*t* − *τ*) and *Y*, which is unique with respect to the past of the receiver *X*_2_ (*t* − *τ*):

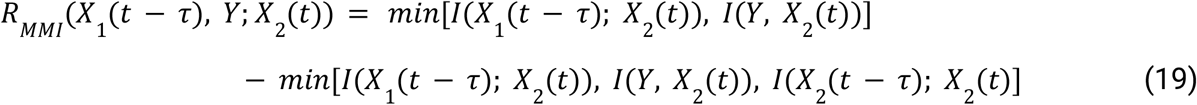

Therefore, the FIT measure is defined as the minimum between these two atoms (Eq. 14 and 15) to select the smallest nonnegative piece of information:

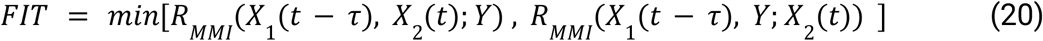

We computed the FIT measure between all pairs of 8 sets of brain regions at different delays tau ranging from −0.005 to −0.2s, and averaged over delays. Given that the scope of the study was not a detailed characterisation of time delays between clusters, we used such averaging procedure, as in previous papers (Celotto et al, 2023). Statistical analysis was similarly applied as for all other measures.

### Statistical analysis

For the statistical inferences, we used a group-level approach based on non-parametric permutations and encompassing non-negative measures of information developed in a previous study (Combrisson, Allegra, et al. 2022). We used a random effect (RFX) to take into account the inter-subject variability. In this approach, the information theoretical metrics (e.g., mutual information, MI) between the neurophysiological signal and the behavioural regressor are computed across trials for each participant separately, at each time point and brain region. In order to assess whether the estimated effect size in MI significantly differed from the chance distribution and to correct for multiple comparisons, we implemented a cluster-wise statistics approach (Maris and Oostenveld, 2007). For cluster-level statistics, the cluster forming threshold is defined as the 95th percentile across all of the permutations. Such threshold is used to identify the clusters on both the true and permuted data. To sample the distribution of MI attainable by chance, we computed the MI between the brain data and a randomly shuffled version of the behavioural variable. This procedure was then repeated 1000 times. We took the mean of the MI values computed on the permutations, and used this mean (MI) to perform a one sample t-test across all the participants’ MI values obtained both from original and permuted data. We then used a cluster-based approach to assess whether the size of the estimated t-values significantly differs from its distribution. The cluster forming threshold was defined as the 95th percentile of the distribution of t-values. We used this cluster forming threshold to identify the cluster mass of t-values on both original and permuted data. As a reminder, the cluster-based approach (Maris and Oostenveld, 2007) is performed on the mass of the clusters (i.e., the summed activity above the threshold), rather than on individual temporal samples. Thus, the p-value of the cluster is not representative of the individual samples within the cluster and cannot be interpreted as a measure of the onset and offset of significant effects. Finally, to correct for multiple comparisons across both time and space, we build a distribution made of the 1000 largest clusters estimated on the permuted data. The final corrected p-values were inferred as the proportion of permutations exceeding the t-values. For what concerns functional connectivity analysis, the information theoretical metrics of interest (e.g., redundant connectivity) across pairs of brain regions were studied similarly as the local MI time courses.

## Acknowledgements

A.B., M.J., M.K. and E.C were supported by the PRC project “CausaL” (ANR-18-CE28-0016). A.B. was supported by A*Midex Foundation of Aix-Marseille University project “Hinteract” (AMX-22-RE-AB-071). R.B. has received funding from the French government under the “France 2030” investment plan managed by the French National Research Agency (reference : ANR-16-CONV000X /ANR-17-EURE-0029) and from Excellence Initiative of AixMarseille University - A*MIDEX (AMX-19-IET-004). The “Center de Calcul Intensif of the Aix-Marseille University (CCIAM)” is acknowledged for high-performance computing resources. A.B and E.C were also supported by EU’s Horizon 2020 Framework Programme for Research and Innovation under the Specific Grant Agreements No. 101147319 (EBRAINS 2.0 Project) and No. 945539 (Human Brain Project SGA3). We thank Jean-Michel Badier and the MEG center of Aix Marseille University (INSERM, INS, Inst Neurosci Syst, Marseille, France) for support in MEG experiments.

## Code and data availability

Higher-order redundancy and synergistic measures were computed using the HOI toolbox (https://github.com/brainets/hoi). All pair-wise information theoretical measures, including the FIT, and statistical analyses were performed using the Frites Python package (https://github.com/brainets/frites) (Combrisson, Basanisi, et al. 2022).

## REFERENCES

Adams, Rick A., Stewart Shipp, and Karl J. Friston. 2013. “Predictions Not Commands: Active Inference in the Motor System.” Brain Structure & Function 218 (3): 611–643.

Allan, Lorraine G., Samuel D. Hannah, Matthew J. C. Crump, and Shepard Siegel. 2008. “The Psychophysics of Contingency Assessment.” Journal of Experimental Psychology. General 137 (2): 226–243.

Averbeck, Bruno B., and Vincent D. Costa. 2017. “Motivational Neural Circuits Underlying Reinforcement Learning.” Nature Neuroscience 20 (4): 505–512.

Averbeck, Bruno, and John P. O’Doherty. 2021. “Reinforcement-Learning in Fronto-Striatal Circuits.” Neuropsychopharmacology: Official Publication of the American College of Neuropsychopharmacology 47 (1): 147–162.

Badre, David, Bradley B. Doll, Nicole M. Long, and Michael J. Frank. 2012. “Rostrolateral Prefrontal Cortex and Individual Differences in Uncertainty-Driven Exploration.” Neuron 73 (3): 595–607.

Baldi, Pierre, and Laurent Itti. 2010. “Of Bits and Wows: A Bayesian Theory of Surprise with Applications to Attention.” Neural Networks: The Official Journal of the International Neural Network Society 23 (5): 649–666.

Ballesta, Sébastien, Weikang Shi, Katherine E. Conen, and Camillo Padoa-Schioppa. 2020. “Values Encoded in Orbitofrontal Cortex Are Causally Related to Economic Choices.” Nature 588 (7838): 450–453.

Bandura, Albert. 1997. Self-Efficacy: The Exercise of Control. (New York, NY, US) 604. https://psycnet.apa.org/fulltext/1997-08589-000.pdf.

Barrett, Adam B. 2015. “Exploration of Synergistic and Redundant Information Sharing in Static and Dynamical Gaussian Systems.” Physical Review. E, Statistical, Nonlinear, and Soft Matter Physics 91 (5): 052802.

Bartolo, Ramon, and Bruno B. Averbeck. 2020. “Prefrontal Cortex Predicts State Switches during Reversal Learning.” Neuron 106 (6): 1044–1054.e4.

Bressler, Steven L., and Anil K. Seth. 2011. “Wiener–Granger Causality: A Well Established Methodology.” NeuroImage 58 (2): 323–329.

Bromberg-Martin, Ethan S., Yang-Yang Feng, Takaya Ogasawara, J. Kael White, Kaining Zhang, and Ilya E. Monosov. 2024. “A Neural Mechanism for Conserved Value Computations Integrating Information and Rewards.” Nature Neuroscience 27 (1): 159–175.

Bromberg-Martin, Ethan S., and Okihide Hikosaka. 2009. “Midbrain Dopamine Neurons Signal Preference for Advance Information about Upcoming Rewards.” Neuron 63 (1): 119–126.

Bromberg-Martin, Ethan S., and Okihide Hikosaka. 2011. “Lateral Habenula Neurons Signal Errors in the Prediction of Reward Information.” Nature Neuroscience 14 (9): 1209–1216.

Brovelli, Andrea, Jean-Michel Badier, Francesca Bonini, Fabrice Bartolomei, Olivier Coulon, and Guillaume Auzias. 2017. “Dynamic Reconfiguration of Visuomotor-Related Functional Connectivity Networks.” The Journal of Neuroscience : The Official Journal of the Society for Neuroscience 37 (4): 839–853.

Brovelli, Andrea, Daniel Chicharro, Jean-Michel Badier, Huifang Wang, and Viktor Jirsa. 2015. “Characterization of Cortical Networks and Corticocortical Functional Connectivity Mediating Arbitrary Visuomotor Mapping.” The Journal of Neuroscience : The Official Journal of the Society for Neuroscience 35 (37): 12643–12658.

Brovelli, Andrea, Mingzhou Ding, Anders Ledberg, Yonghong Chen, Richard Nakamura, and Steven L. Bressler. 2004. “Beta Oscillations in a Large-Scale Sensorimotor Cortical Network: Directional Influences Revealed by Granger Causality.” Proceedings of the National Academy of Sciences of the United States of America 101 (26): 9849–9854.

Brovelli, Andrea, Jean-Philippe Lachaux, Philippe Kahane, and Driss Boussaoud. 2005. “High Gamma Frequency Oscillatory Activity Dissociates Attention from Intention in the Human Premotor Cortex.” NeuroImage 28 (1): 154–164.

Brydevall, Maja, Daniel Bennett, Carsten Murawski, and Stefan Bode. 2018. “The Neural Encoding of Information Prediction Errors during Non-Instrumental Information Seeking.” Scientific Reports 8 (1): 6134.

Cai, Xinying, and Camillo Padoa-Schioppa. 2014. “Contributions of Orbitofrontal and Lateral Prefrontal Cortices to Economic Choice and the Good-to-Action Transformation.” Neuron 81 (5): 1140–1151.

Celotto, Marco, Jan Bím, Alejandro Tlaie, et al. 2023. “An Information-Theoretic Quantification of the Content of Communication between Brain Regions.” bioRxiv : The Preprint Server for Biology, ahead of print, June 14. 10.1101/2023.06.14.544903.

Charpentier, Caroline J., Ethan S. Bromberg-Martin, and Tali Sharot. 2018. “Valuation of Knowledge and Ignorance in Mesolimbic Reward Circuitry.” Proceedings of the National Academy of Sciences of the United States of America 115 (31): E7255–E7264.

Cockburn, Jeffrey, Vincent Man, William A. Cunningham, and John P. O’Doherty. 2022. “Novelty and Uncertainty Regulate the Balance between Exploration and Exploitation through Distinct Mechanisms in the Human Brain.” Neuron 110 (16): 2691–2702.e8.

Cohen, Jonathan D., Samuel M. McClure, and Angela J. Yu. 2007. “Should I Stay or Should I Go? How the Human Brain Manages the Trade-off between Exploitation and Exploration.” Philosophical Transactions of the Royal Society of London. Series B, Biological Sciences 362 (1481): 933–942.

Combrisson, Etienne, Michele Allegra, Ruggero Basanisi, et al. 2022. “Group-Level Inference of Information-Based Measures for the Analyses of Cognitive Brain Networks from Neurophysiological Data.” NeuroImage 258 (September): 119347.

Combrisson, Etienne, Ruggero Basanisi, Vinicius Lima Cordeiro, Robin A. A. Ince, and Andrea Brovelli. 2022. “Frites: A Python Package for Functional Connectivity Analysis and Group-Level Statistics of Neurophysiological Data.” Journal of Open Source Software 7 (79): 3842.

Combrisson, Etienne, Ruggero Basanisi, Maelle C. M. Gueguen, et al. 2024. Neural Interactions in the Human Frontal Cortex Dissociate Reward and Punishment Learning. June 28. 10.7554/eLife.92938.

Combrisson, Etienne, Ruggero Basanisi, Matteo Neri, et al. 2025. “Higher-Order and Distributed Synergistic Functional Interactions Encode Information Gain in Goal-Directed Learning.” Nature Communications 16 (1): 7179.

Coupe, P., P. Yger, S. Prima, P. Hellier, C. Kervrann, and C. Barillot. 2008. “An Optimized Blockwise Nonlocal Means Denoising Filter for 3-D Magnetic Resonance Images.” IEEE Transactions on Medical Imaging 27 (4): 425–441.

Crone, Nathan E., Alon Sinai, and Anna Korzeniewska. 2006. “High-Frequency Gamma Oscillations and Human Brain Mapping with Electrocorticography.” Progress in Brain Research 159: 275–295.

Daw, Nathaniel D., and John P. O’Doherty. 2014. “Multiple Systems for Value Learning.” In Neuroeconomics. Elsevier.

Daw, Nathaniel D., John P. O’Doherty, Peter Dayan, Ben Seymour, and Raymond J. Dolan. 2006. “Cortical Substrates for Exploratory Decisions in Humans.” Nature 441 (7095): 876–879.

Destrieux, Christophe, Bruce Fischl, Anders Dale, and Eric Halgren. 2010. “Automatic Parcellation of Human Cortical Gyri and Sulci Using Standard Anatomical Nomenclature.” NeuroImage 53 (1): 1–15.

Dickinson, Anthony. 1994. “CHAPTER 3 - Instrumental Conditioning.” In Animal Learning and Cognition, edited by N. J. Mackintosh. Academic Press.

Dolan, Ray J., and Peter Dayan. 2013. “Goals and Habits in the Brain.” Neuron 80 (2): 312–325.

Faraji, Mohammadjavad, Kerstin Preuschoff, and Wulfram Gerstner. 2018. “Balancing New against Old Information: The Role of Puzzlement Surprise in Learning.” Neural Computation 30 (1): 34–83.

Findling, Charles, Vasilisa Skvortsova, Rémi Dromnelle, Stefano Palminteri, and Valentin Wyart. 2019. “Computational Noise in Reward-Guided Learning Drives Behavioral Variability in Volatile Environments.” Nature Neuroscience 22 (12): 2066–2077.

Findling, Charles, and Valentin Wyart. 2024. “Computation Noise Promotes Zero-Shot Adaptation to Uncertainty during Decision-Making in Artificial Neural Networks.” Science Advances 10 (44): eadl3931.

Fouragnan, Elsa, Chris Retzler, and Marios G. Philiastides. 2018. “Separate Neural Representations of Prediction Error Valence and Surprise: Evidence from an fMRI Meta-Analysis.” Human Brain Mapping 39 (7): 2887–2906.

Frank, Michael J., Bradley B. Doll, Jen Oas-Terpstra, and Francisco Moreno. 2009. “Prefrontal and Striatal Dopaminergic Genes Predict Individual Differences in Exploration and Exploitation.” Nature Neuroscience 12 (8): 1062–1068.

Friston, Karl, Thomas FitzGerald, Francesco Rigoli, Philipp Schwartenbeck, and Giovanni Pezzulo. 2017. “Active Inference: A Process Theory.” Neural Computation 29 (1): 1–49.

Friston, Karl J., Jean Daunizeau, James Kilner, and Stefan J. Kiebel. 2010. “Action and Behavior: A Free-Energy Formulation.” Biological Cybernetics 102 (3): 227–260.

Friston, Karl, Francesco Rigoli, Dimitri Ognibene, Christoph Mathys, Thomas Fitzgerald, and Giovanni Pezzulo. 2015. “Active Inference and Epistemic Value.” Cognitive Neuroscience 6 (4): 187–214.

Gardner, Matthew P. H., Davied Sanchez, Jessica C. Conroy, Andrew M. Wikenheiser, Jingfeng Zhou, and Geoffrey Schoenbaum. 2020. “Processing in Lateral Orbitofrontal Cortex Is Required to Estimate Subjective Preference during Initial, but Not Established, Economic Choice.” Neuron 108 (3): 526–537.e4.

Gershman, Samuel J. 2018. “Deconstructing the Human Algorithms for Exploration.” Cognition 173 (April): 34–42.

Gläscher, Jan, Nathaniel Daw, Peter Dayan, and John P. O’Doherty. 2010. “States versus Rewards: Dissociable Neural Prediction Error Signals Underlying Model-Based and Model-Free Reinforcement Learning.” Neuron 66 (4): 585–595.

Goddu, Mariel K., and Alison Gopnik. 2024. “The Development of Human Causal Learning and Reasoning.” Nature Reviews Psychology 3 (5): 319–339.

Gore, Felicity, Melissa Hernandez, Charu Ramakrishnan, Ailey K. Crow, Robert C. Malenka, and Karl Deisseroth. 2023. “Orbitofrontal Cortex Control of Striatum Leads Economic Decision-Making.” Nature Neuroscience 26 (9): 1566–1574.

Gottlieb, Jacqueline, and Pierre-Yves Oudeyer. 2018. “Towards a Neuroscience of Active Sampling and Curiosity.” Nature Reviews. Neuroscience 19 (12): 758–770.

Gottlieb, Jacqueline, Pierre-Yves Oudeyer, Manuel Lopes, and Adrien Baranes. 2013. “Information-Seeking, Curiosity, and Attention: Computational and Neural Mechanisms.” Trends in Cognitive Sciences 17 (11): 585–593.

Gramfort, Alexandre, Martin Luessi, Eric Larson, et al. 2013. “MEG and EEG Data Analysis with MNE-Python.” Frontiers in Neuroscience 7 (December): 267.

Granger, C. W. J. 1980. “Testing for Causality: A Personal Viewpoint.” Journal of Economic Dynamics & Control 2 (January): 329–352.

Gross, J., J. Kujala, M. Hamalainen, L. Timmermann, A. Schnitzler, and R. Salmelin. 2001. “Dynamic Imaging of Coherent Sources: Studying Neural Interactions in the Human Brain.” Proceedings of the National Academy of Sciences of the United States of America 98 (2): 694–699.

Haber, Suzanne N., and Brian Knutson. 2009. “The Reward Circuit: Linking Primate Anatomy and Human Imaging.” Neuropsychopharmacology: Official Publication of the American College of Neuropsychopharmacology 35 (1): 4–26.

Hafner, Danijar, Jurgis Pasukonis, Jimmy Ba, and Timothy Lillicrap. 2025. “Mastering Diverse Control Tasks through World Models.” Nature 640 (8059): 647–653.

Ince, Robin A. A., Bruno L. Giordano, Christoph Kayser, Guillaume A. Rousselet, Joachim Gross, and Philippe G. Schyns. 2017. “A Statistical Framework for Neuroimaging Data Analysis Based on Mutual Information Estimated via a Gaussian Copula.” Human Brain Mapping 38 (3): 1541–1573.

Itti, Laurent, and Pierre Baldi. 2009. “Bayesian Surprise Attracts Human Attention.” Vision Research 49 (10): 1295–1306.

Jas, Mainak, Denis A. Engemann, Yousra Bekhti, Federico Raimondo, and Alexandre Gramfort. 2017. “Autoreject: Automated Artifact Rejection for MEG and EEG Data.” NeuroImage 159 (October): 417–429.

Jenkins, H. M., and W. C. Ward. 1965. “JUDGMENT OF CONTINGENCY BETWEEN RESPONSES AND OUTCOMES.” Psychological Monographs 79: SUPPL 1:1–17.

Jerbi, Karim, Tomás Ossandón, Carlos M. Hamamé, et al. 2009. “Task-Related Gamma-Band Dynamics from an Intracerebral Perspective: Review and Implications for Surface EEG and MEG.” Human Brain Mapping 30 (6): 1758–1771.

Jocham, Gerhard, Kay H. Brodersen, Alexandra O. Constantinescu, et al. 2016. “Reward-Guided Learning with and without Causal Attribution.” Neuron 90 (1): 177–190.

Khamassi, Mehdi, Marceau Nahon, and Raja Chatila. 2024. “Strong and Weak Alignment of Large Language Models with Human Values.” Scientific Reports 14 (1): 19399.

Kidd, Celeste, and Benjamin Y. Hayden. 2015. “The Psychology and Neuroscience of Curiosity.” Neuron 88 (3): 449–460.

Kobayashi, Kenji, and Ming Hsu. 2019. “Common Neural Code for Reward and Information Value.” Proceedings of the National Academy of Sciences of the United States of America 116 (26): 13061–13066.

Kobayashi, Kenji, and Joseph W. Kable. 2024. “Neural Mechanisms of Information Seeking.” Neuron 112 (11): 1741–1756.

Kobayashi, Kenji, Sangil Lee, Alexandre L. S. Filipowicz, Kara D. McGaughey, Joseph W. Kable, and Matthew R. Nassar. 2021. “Dynamic Representation of the Subjective Value of Information.” The Journal of Neuroscience: The Official Journal of the Society for Neuroscience 41 (39): 8220–8232.

Lachaux, Jean-Philippe, Pierre Fonlupt, Philippe Kahane, et al. 2007. “Relationship between Task-Related Gamma Oscillations and BOLD Signal: New Insights from Combined fMRI and Intracranial EEG.” Human Brain Mapping 28 (12): 1368–1375.

Lee, Sang Wan, Shinsuke Shimojo, and John P. O’Doherty. 2014. “Neural Computations Underlying Arbitration between Model-Based and Model-Free Learning.” Neuron 81 (3): 687–699.

Levy, Dino J., and Paul W. Glimcher. 2012. “The Root of All Value: A Neural Common Currency for Choice.” Current Opinion in Neurobiology 22 (6): 1027–1038.

Liakoni, Vasiliki, Marco P. Lehmann, Alireza Modirshanechi, et al. 2022. “Brain Signals of a Surprise-Actor-Critic Model: Evidence for Multiple Learning Modules in Human Decision Making.” NeuroImage 246 (February): 118780.

Liakoni, Vasiliki, Alireza Modirshanechi, Wulfram Gerstner, and Johanni Brea. 2021. “Learning in Volatile Environments With the Bayes Factor Surprise.” Neural Computation 33 (2): 269–340.

Liljeholm, Mimi, Elizabeth Tricomi, John P. O’Doherty, and Bernard W. Balleine. 2011. “Neural Correlates of Instrumental Contingency Learning: Differential Effects of Action-Reward Conjunction and Disjunction.” The Journal of Neuroscience : The Official Journal of the Society for Neuroscience 31 (7): 2474–2480.

Liljeholm, Mimi, Shuo Wang, June Zhang, and John P. O’Doherty. 2013. “Neural Correlates of the Divergence of Instrumental Probability Distributions.” The Journal of Neuroscience: The Official Journal of the Society for Neuroscience 33 (30): 12519–12527.

Lizier, Joseph T., Nils Bertschinger, Jürgen Jost, and Michael Wibral. 2018. “Information Decomposition of Target Effects from Multi-Source Interactions: Perspectives on Previous, Current and Future Work.” Entropy 20 (4). 10.3390/e20040307.

Luppi, Andrea I., Pedro A. M. Mediano, Fernando E. Rosas, et al. 2022. “A Synergistic Core for Human Brain Evolution and Cognition.” Nature Neuroscience 25 (6): 771–782.

Mehlhorn, Katja, Ben R. Newell, Peter M. Todd, et al. 2015. “Unpacking the Exploration–exploitation Tradeoff: A Synthesis of Human and Animal Literatures.” Decisions 2 (3): 191–215.

Miller, E. K., and J. D. Cohen. 2001. “An Integrative Theory of Prefrontal Cortex Function.” Annual Review of Neuroscience 24: 167–202.

Modirshanechi, Alireza, Sophia Becker, Johanni Brea, and Wulfram Gerstner. 2023. “Surprise and Novelty in the Brain.” Current Opinion in Neurobiology 82 (October): 102758.

Modirshanechi, Alireza, Johanni Brea, and Wulfram Gerstner. 2022. “A Taxonomy of Surprise Definitions.” Journal of Mathematical Psychology 110 (September): 102712.

Modirshanechi, Alireza, Wei-Hsiang Lin, He A. Xu, Michael H. Herzog, and Wulfram Gerstner. 2025. “Novelty as a Drive of Human Exploration in Complex Stochastic Environments.” Proceedings of the National Academy of Sciences of the United States of America 122 (39): e2502193122.

Morris, Richard W., Amir Dezfouli, Kristi R. Griffiths, and Bernard W. Balleine. 2014. “Action-Value Comparisons in the Dorsolateral Prefrontal Cortex Control Choice between Goal-Directed Actions.” Nature Communications 5 (July): 4390.

Morris, Richard W., Amir Dezfouli, Kristi R. Griffiths, Mike E. Le Pelley, and Bernard W. Balleine. 2022. “The Neural Bases of Action-Outcome Learning in Humans.” The Journal of Neuroscience: The Official Journal of the Society for Neuroscience, ahead of print, March 16. 10.1523/JNEUROSCI.1079-21.2022.

Mukamel, Roy, Hagar Gelbard, Amos Arieli, Uri Hasson, Itzhak Fried, and Rafael Malach. 2005. “Coupling between Neuronal Firing, Field Potentials, and FMRI in Human Auditory Cortex.” Science (New York, N.Y.) 309 (5736): 951–954.

Niessing, Jörn, Boris Ebisch, Kerstin E. Schmidt, Michael Niessing, Wolf Singer, and Ralf A. W. Galuske. 2005. “Hemodynamic Signals Correlate Tightly with Synchronized Gamma Oscillations.” Science (New York, N.Y.) 309 (5736): 948–951.

Nir, Yuval, Lior Fisch, Roy Mukamel, et al. 2007. “Coupling between Neuronal Firing Rate, Gamma LFP, and BOLD fMRI Is Related to Interneuronal Correlations.” Current Biology : CB 17 (15): 1275–1285.

Norton, Kaitlyn G., and Mimi Liljeholm. 2020. “The Rostrolateral Prefrontal Cortex Mediates a Preference for High-Agency Environments.” The Journal of Neuroscience: The Official Journal of the Society for Neuroscience 40 (22): 4401–4409.

“OSF.” n.d. Accessed May 7, 2026. 10.31234/osf.io/du3e2_v1.

Padoa-Schioppa, Camillo, and John A. Assad. 2006. “Neurons in the Orbitofrontal Cortex Encode Economic Value.” Nature 441 (7090): 223–226.

Parr, Thomas, Giovanni Pezzulo, and Karl J. Friston. 2022. Active Inference: The Free Energy Principle in Mind, Brain, and Behavior. MIT Press.

Pearl, Judea, and Dana Mackenzie. 2018. The Book of Why: The New Science of Cause and Effect. Penguin UK.

Percival, Donald B., and Andrew T. Walden. 1993. Spectral Analysis for Physical Applications. Cambridge University Press.

Pezzulo, Giovanni, Thomas Parr, Paul Cisek, Andy Clark, and Karl Friston. 2024. “Generating Meaning: Active Inference and the Scope and Limits of Passive AI.” Trends in Cognitive Sciences 28 (2): 97–112.

Rangel, Antonio, Colin Camerer, and P. Read Montague. 2008. “A Framework for Studying the Neurobiology of Value-Based Decision Making.” Nature Reviews. Neuroscience 9 (7): 545–556.

Ray, Supratim, and John H. R. Maunsell. 2011. “Different Origins of Gamma Rhythm and High-Gamma Activity in Macaque Visual Cortex.” PLOS Biology 9 (4): e1000610.

Rushworth, Matthew F. S., Maryann P. Noonan, Erie D. Boorman, Mark E. Walton, and Timothy E. Behrens. 2011. “Frontal Cortex and Reward-Guided Learning and Decision-Making.” Neuron 70 (6): 1054–1069.

Schreiber, T. 2000. “Measuring Information Transfer.” Physical Review Letters 85 (2): 461–464.

Schwartenbeck, Philipp, Johannes Passecker, Tobias U. Hauser, Thomas Hb FitzGerald, Martin Kronbichler, and Karl J. Friston. 2019. “Computational Mechanisms of Curiosity and Goal-Directed Exploration.” eLife 8 (May). 10.7554/eLife.41703.

Shanks, D. R., and A. Dickinson. 1991. “Instrumental Judgment and Performance under Variations in Action-Outcome Contingency and Contiguity.” Memory & Cognition 19 (4): 353–360.

Shipp, Stewart, Rick A. Adams, and Karl J. Friston. 2013. “Reflections on Agranular Architecture: Predictive Coding in the Motor Cortex.” Trends in Neurosciences 36 (12): 706–716.

Sutton, Richard S., and Andrew G. Barto. 2018. Reinforcement Learning, Second Edition: An Introduction. MIT Press.

Tanaka, Saori C., Bernard W. Balleine, and John P. O’Doherty. 2008. “Calculating Consequences: Brain Systems That Encode the Causal Effects of Actions.” The Journal of Neuroscience: The Official Journal of the Society for Neuroscience 28 (26): 6750–6755.

Tishby, Naftali, and Daniel Polani. 2011. “Information Theory of Decisions and Actions.” In Perception-Action Cycle: Models, Architectures, and Hardware, edited by Vassilis Cutsuridis, Amir Hussain, and John G. Taylor. Springer New York.

Tolman, E. C. 1948. “Cognitive Maps in Rats and Men.” Psychological Review 55 (4): 189–208.

Walton, Mark E., Timothy E. J. Behrens, Mark J. Buckley, Peter H. Rudebeck, and Matthew F. S. Rushworth. 2010. “Separable Learning Systems in the Macaque Brain and the Role of Orbitofrontal Cortex in Contingent Learning.” Neuron 65 (6): 927–939.

Wang, Huifang E., Julia Scholly, Paul Triebkorn, et al. 2021. “VEP Atlas: An Anatomic and Functional Human Brain Atlas Dedicated to Epilepsy Patients.” Journal of Neuroscience Methods 348 (January): 108983.

“Website.” n.d. 10.1037/h0081013.

White, J. Kael, Ethan S. Bromberg-Martin, Sarah R. Heilbronner, et al. 2019. “A Neural Network for Information Seeking.” Nature Communications 10 (1): 5168.

Williams, Paul L., and Randall D. Beer. 2010. “Nonnegative Decomposition of Multivariate Information.” In arXiv [cs.IT]. April 14. arXiv. http://arxiv.org/abs/1004.2515.

Wilson, Robert C., Andra Geana, John M. White, Elliot A. Ludvig, and Jonathan D. Cohen. 2014. “Humans Use Directed and Random Exploration to Solve the Explore-Exploit Dilemma.” Journal of Experimental Psychology. General 143 (6): 2074–2081.

Yu, Angela J., and Peter Dayan. 2005. “Uncertainty, Neuromodulation, and Attention.” Neuron 46 (4): 681–692.

